# Fast, Accurate, and Versatile Data Analysis Platform for the Quantification of Molecular Spatiotemporal Signals

**DOI:** 10.1101/2024.05.02.592259

**Authors:** Xuelong Mi, Alex Bo-Yuan Chen, Daniela Duarte, Erin Carey, Charlotte R. Taylor, Philipp N. Braaker, Mark Bright, Rafael G. Almeida, Jing-Xuan Lim, Virginia M. S. Ruetten, Wei Zheng, Mengfan Wang, Michael E. Reitman, Yizhi Wang, Kira E. Poskanzer, David A. Lyons, Axel Nimmerjahn, Misha B. Ahrens, Guoqiang Yu

**Author notes:** These authors contributed equally (Xuelong Mi, Alex Bo-Yuan Chen).

## Abstract

Optical recording of intricate molecular dynamics is becoming an indispensable technique for biological studies, accelerated by the development of new or improved biosensors and microscopy technology. This creates major computational challenges to extract and quantify biologically meaningful spatiotemporal patterns embedded within complex and rich data sources, many of which cannot be captured with existing methods. Here, we introduce Activity Quantification and Analysis (AQuA2), a fast, accurate, and versatile data analysis platform built upon advanced machine learning techniques. It decomposes complex live imaging-based datasets into elementary signaling events, allowing accurate and unbiased quantification of molecular activities and identification of consensus functional units. We demonstrate applications across a wide range of biosensors, cell types, organs, animal models, and imaging modalities. As exemplar findings, we show how AQuA2 identified drug-dependent interactions between neurons and astroglia, and distinct sensorimotor signal propagation patterns in the mouse spinal cord.

## INTRODUCTION

Imaging cellular and molecular activity across space and time has emerged as a crucial approach in many fields, such as neuroscience ^1^, cell biology ^2^, pathology ^3^, and developmental biology ^4^. Recent developments in modern genetically encoded fluorescent probes ^5–7^ and advanced imaging techniques ^8–10^ have enabled the observation of a wide range of signals, including calcium ions, ATP, neurotransmitters, neuromodulators, and other molecules, greatly expanding the breadth and depth of scientific studies. However, with the rapid growth of data generation and the revealing of complex spatiotemporal activity patterns such as heterogeneous spatial footprints and various propagations (see top row of Figure 3), quantifying and understanding the data has become a limiting factor. Manual inspection is simply infeasible due to the complexity and large data size, and even when possible, it often misses subtle yet important information. Although automated image analysis methods have been developed, they are typically constrained to modeling a specific type of signal with an assumption of simple spatiotemporal pattern or plagued by low accuracy, extensive processing times, and a limited set of analysis functions, failing to meet today’s demands for a unified data analysis platform that allows flexible and accurate quantification of diverse and complex data from various experimental settings.

Existing fluorescent imaging analysis techniques can be broadly classified into two categories: region-of-interest-based (ROI-based) methods ^11–13^ and event-based methods ^14–16^. ROI-based methods, such as suite2p ^12^ and CaImAn ^11^, rely on identifying regions of interest (ROIs), which are ill-suited to capture the spatial dynamics of molecular signals. An ROI is a fixed spatial area associated with a single temporal dynamic. While certain ROI methods, such as non-negative matrix factorization (NMF)-based approaches ^11,17^, permit overlap between regions, they are based on weighted averages of a low number of temporal components and do not allow the spatial dynamics to be faithfully captured. These methods can be effective for analyzing stereotypical neuronal signals. However, their stringent assumption of spatial stationarity often results in suboptimal performance and frequently distorted descriptions when the target signals exhibit complex and flexible spatiotemporal dynamics. Event-based methods, in contrast, are specifically designed to capture dynamic activities with intricate spatiotemporal features. Astrocyte Quantitative Analysis (AQuA) ^14^, developed by our team and widely used in the astrocyte field, pioneered event-based quantification. Because events are jointly determined by the spatial coherence and the temporal pattern, it allows for more flexible modeling of both spatial and temporal dynamics than traditional ROI-based models, in that one pixel can participate in different kinds of events, and any one signal can propagate across space. This makes AQuA widely adopted for quantifying astrocyte calcium signals that can flow along the geometry of the cell and exhibit subcellular and population-wide spatial dynamics.

With growing AQuA usage^18–21^, we, as its developer, have received numerous requests for a fast, accurate, and versatile platform with more functions to enhance the quantification and analysis of generic molecular spatiotemporal activity. First, researchers find region or location information helpful in interpreting their results, and therefore, they need analysis methods that integrate region-based and event-based approaches. Second, as multiplexing imaging methods are maturing ^22,23^, there is a need to model more than one type of signal and analyze the interactions between them. Third, since scientists aim to minimize the disturbance to the system under investigation, data sets with low signal-to-noise ratios are generated, requiring more accurate algorithms to cope with large noise. Fourth, new data is getting larger and larger, frequently involving three spatial dimensions. The large data size necessitates better approaches in terms of both computational time and computer memory. Finally, although AQuA was originally designed primarily for astrocyte calcium activity, it has been applied to many other cell types and signals without thorough validation and optimization.

Responding to the feedback and requests, we developed Activity Quantification and Analysis (AQuA2), an entirely new version of AQuA. Now, the first letter A in AQuA2 stands for activity instead of astrocyte as in AQuA, indicating the great expansion of applicability. We introduce the Consensus Functional Unit (CFU) concept to integrate the ROI-based and event-based approaches. Potential functional units, termed CFUs, are detected with the hypothesis that a functional unit is expected to exhibit multiple events with a consistent spatial footprint/region. The CFU concept, bridging region and event definitions, can be considered a more flexible version of ROI, allowing signals to have different sizes, shapes, and propagation patterns while maintaining consistent spatial foundations. This new function can facilitate deeper biological insights by utilizing strengths from both ROI-based and event-based approaches. Moreover, introducing CFUs enables us to analyze the interaction between signals recorded by different biosensors. To improve the accuracy when used on noisy data, we designed a more reliable top-down strategy, which utilizes more information from a broader field of view than the original bottom-up approach. This new strategy integrates principles of probability theory while adopting innovative machine learning algorithms such as BIdirectional pushing with Linear Component Operations (BILCO) ^24^ for ever-growing data volumes. It can efficiently and accurately quantify signals with various features. Furthermore, to enhance usability, we have equipped AQuA2 with a user-friendly interface compatible with processing 2D, 3D, and dual-color data.

AQuA2 surpasses existing ROI- and event-based methods (suite2p ^12^, CaImAn ^11^, Begonia ^16^, and AQuA ^14^) in accuracy and demonstrates high efficiency, corroborated through extensive testing on both simulated and real-world data. Many additional signal properties, such as the potential functional units, the interaction between signals, and 3D signal propagation, can be readily captured by AQuA2 but difficult or impossible with existing methods, individually or combined. The user interface is optimized so that the operation is intuitive and detection parameters can be set based on known biological constraints. Utilizing AQuA2, we quantified the dynamics of multiple distinct biological signals (calcium, norepinephrine (NE) ^5^, dopamine ^25^, ATP ^6^, and acetylcholine (Ach) ^26^), cell types (neurons, astrocytes, microglia, and oligodendrocytes), organs (brains and spinal cords), animals (mouse and zebrafish), and imaging modalities (confocal, two-photon, light sheet). Additionally, using AQuA2, we explored drug-dependent interactions between neurons and astroglia and discovered different modes of sensorimotor signal propagation in the mouse spinal cord. These results establish AQuA2 as a universally applicable toolkit capable of addressing a wide array of scientific inquiries.

## RESULTS

### Event detection pipeline of AQuA2

The event-based methodology, first introduced in Astrocyte Quantitative Analysis (AQuA) ^14^ provides a unique perspective for modeling spatiotemporal activities. It defined an event as a spatiotemporally connected region characterized by fluorescent dynamics, containing a single peak pattern, and originating from a single source. Adjacent pixels belonging to the same event are allowed to exhibit slight delay or deformation in temporal patterns, enabling the analysis of signals with complex features. In AQuA2, we have retained this concept but enhanced the reliability of the event detection pipeline by adopting a top-down framework that incorporates advanced machine learning algorithms. It obtains the signal events by segmenting regions with significant dynamics in the following steps, as depicted in Figure 1B: (1) Baseline estimation and dF (change in fluorescence) calculation according to the non-negative nature of dynamic signals; (2) Active region detection by applying statistical tests on dF; (3) Temporal segmentation through machine learning and image processing techniques to ensure each segmentation (super event) contains a singular peak pattern. (4) Spatial segmentation based on the signal propagation estimated using joint alignment techniques to ensure each final event originates from a single source. For a more detailed explanation of each step, please refer to STAR Methods.

**Figure 1.**
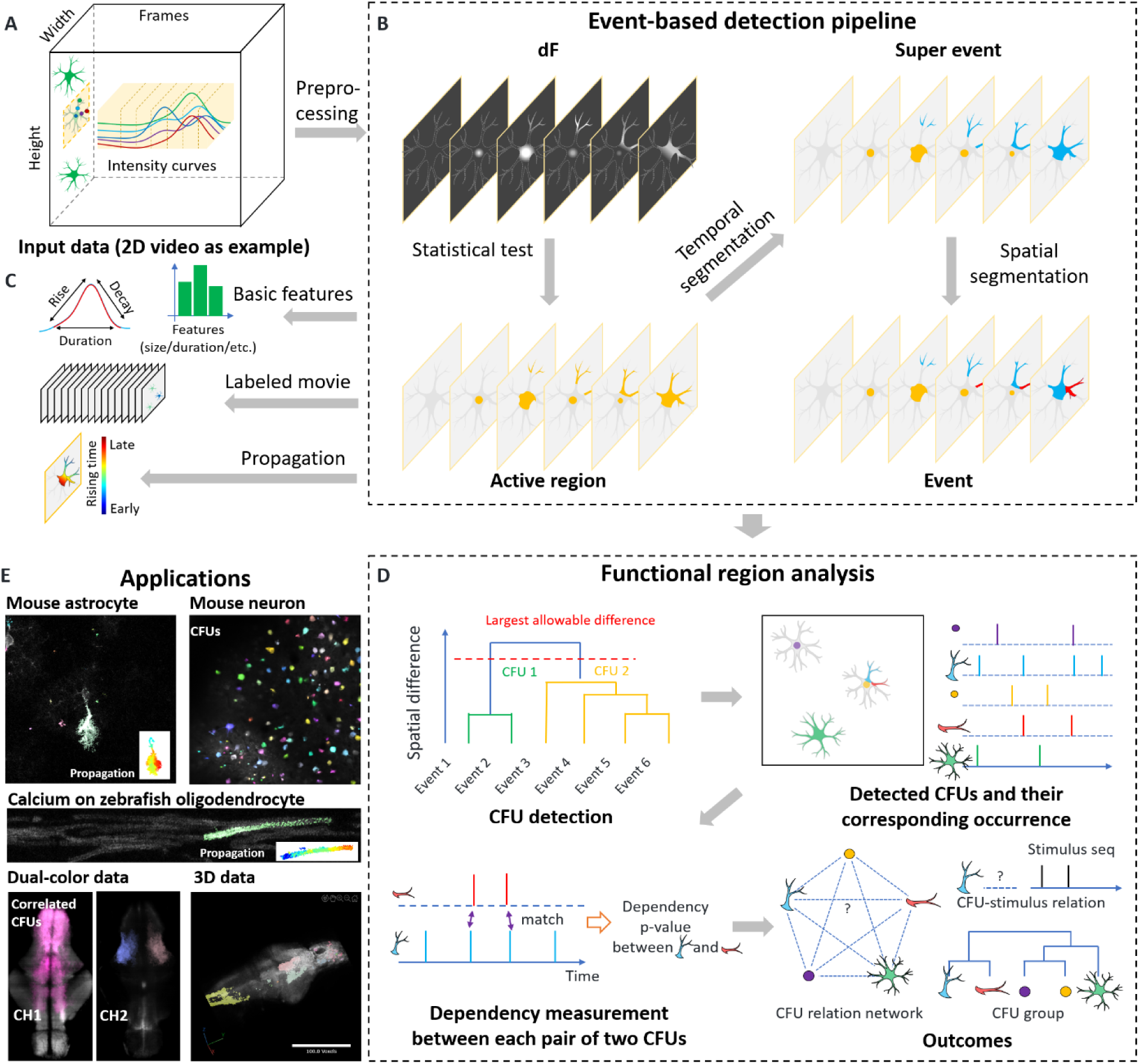
Principles of AQuA2. (A) Typical input data with two spatial dimensions plus a time dimension is shown, with three exemplary cells illustrated in green. The example crop, centered around the middle cell, is utilized in (B), with intensity variations of five positions (labeled by different colors) illustrated by corresponding curves. (B) Event-based signal detection pipeline of AQuA2, including preprocessing, statistical test, temporal segmentation, and spatial segmentation. In the outcome of each step, different colors indicate the corresponding pixels/regions belong to the different detection results. (C) Potential outputs based on event detection pipeline, including extracted event features (size, duration, etc.), movie with event overlay, and propagation map. (D) Diagram for functional region analysis. It consists of Consensus Functional Unit (CFU) detection (cluster events with similar spatial patterns) and interaction analysis between CFUs (take event occurrence sequence as input, output significance p-value of dependency). The outcomes include CFU relation network, CFU-stimulus relation, and CFU groups. (E) Example applications of AQuA2, including signals of different cell types, different biosensors, dual-color data, and 3D data. See also Figure S1.

The new pipeline, in contrast to AQuA, demonstrates superior accuracy and efficiency due to the following key enhancements: Firstly, AQuA2 adopts a top-down framework, which is intrinsically more robust compared to the bottom-up framework used in AQuA. The bottom-up framework, designed to aggregate pixels into events, tends to place excessive emphasis on the temporal information of individual pixels. This makes it susceptible to noise interference, especially in low signal-to-noise ratio (SNR) environments. In contrast, the top-down approach, which identifies signal events by segmenting potential active regions, achieves better performance by integrating the information carried by all the pixels in a broader scope. Combined with the flexibility of event-based methods, this improvement enables AQuA2 to robustly capture a wide range of molecular spatiotemporal activities across various experimental setups. Secondly, by developing and applying advanced machine learning techniques, we made technical innovations in almost every component of the pipeline to achieve accurate analysis with low SNR. These efforts include the development of new methods for baseline fluorescence estimation, noise level estimation, peak detection, integration of prior knowledge, multiscale usage of spatial compactness, split of super-events, and removal of technical artifacts (see STAR Methods). Thirdly, AQuA2 has optimized efficiency by incorporating an innovative machine learning algorithm called BILCO ^24^. This algorithm was specifically designed to address the most computationally intensive module, which accounts for approximately 70% processing time of the whole pipeline and the bottleneck for memory consumption in the original version of AQuA. As a result, there is an average tenfold improvement in both runtime and memory usage in this module. It leads to a more than two-fold acceleration of the entire pipeline compared to AQuA without BILCO, and its efficient memory usage enables the application on ten-fold larger data.

### Identifying consensus functional units and interaction analysis

In response to the limitations of event-based methodologies, which treat individual events as isolated outcomes and are ignorant to the significance of regions with repetitive signals, we introduced the concept of Consensus Functional Units (CFU) and developed the CFU module as an integral component of AQuA2. We hypothesized that if one spatial region generates repeated signal events, it is more likely to be a functional unit, and we refer to such a region as a CFU. This concept offers greater flexibility compared to ROIs, allowing each occurrence of signals to have different sizes, shapes, and propagation patterns while maintaining consistent spatial foundations. On different data types or with different scopes of recordings, CFUs could represent diverse entities, ranging from cellular compartments, individual cells, and cell clusters to tissues and even entire organs.

Based on this assumption, we have employed a hierarchical clustering algorithm to identify CFUs by grouping signals with similar spatial patterns, as shown in Figure 1D. The CFU behind each group can be represented by their average spatial pattern, while the onset time of these signals creates a corresponding event occurrence sequence. This sequence can facilitate the exploration of the interrelationships between CFUs. Compared to the classic ROI-based methods that rely on NMF ^27^ or similar approaches, our CFU identification offers several distinct advantages: Firstly, it inherits the flexibility of the event detection pipeline. Even when signals from the same source exhibit varying sizes, subtle location changes or propagations, our approach can adeptly capture the underlying functional unit. In contrast, ROI-based methods may segment the regions into several arbitrary parts under such conditions (see Figure S4). Secondly, unlike the notoriously hard problem of determining the number of components in ROI-based methods, the CFU module prompts users to set a parameter for the similarity of spatial signal patterns, fostering a more intuitive approach to clustering signals together. Thirdly, CFU identification does not have any bias toward the size of detected components. In contrast, NMF-based methods tend to favor larger components to minimize errors and ignore small functional units.

We have also developed a novel statistical approach for analyzing interactions among identified CFUs. The approach takes sequences of event occurrences in CFUs as its input and assesses the significance of dependency between any pair of two CFUs (see STAR Methods). Compared with traditional methods using correlation, it effectively mitigates the confounding influence of signal shape and duration, enabling precise measurement of interactions among CFUs displaying entirely different signal forms. The interaction analysis, sometimes also called network analysis, is particularly useful when multiple measures are available. For instance, in dual-color neuron and glia data, this approach can discover the associations between neuronal cells and glia, while traditional correlation analysis would fail due to the mismatch of the temporal characteristics of (calcium measurements of) neuronal spikes and glial calcium signals (see Figure 4D). Furthermore, given that stimuli/behavior can be structured as occurrence sequences, this approach can directly investigate the interactions between CFUs and stimuli/behavior. For instance, when recording data from a swimming zebrafish, it can aid in identifying swim-related brain regions (see Figure 6D). Finally, the computed dependency can serve as a measure to cluster spatially separated CFUs into CFU groups, potentially unveiling underlying signaling pathways within the group. For example, when certain brain cells are consistently activated in close temporal proximity, our method can classify these cells together and reveal potential brain circuits through analysis of activation sequences (see Figure S7D).

### AQuA2 is implemented as an open-source MATLAB package, a Fiji Plugin, and a cloud-based web service

We provide AQuA2 software across three distinct environments tailored to various demands: a MATLAB-based package, a Fiji ^28^ plugin written in Java, and a cloud-based online service, AQuA2-Cloud. The MATLAB and Fiji versions allow users to run AQuA2 locally. The MATLAB version makes it easier for users to extend AQuA2, while the Fiji version integrates well into the ImageJ ecosystem. The AQuA2-Cloud enables the offloading of computational tasks to a remote server, which is useful for users without sufficient local computational power for analyzing larger data. This empowers users with lightweight workstations to handle large-scale data through online analysis remotely. All these versions implement the AQuA2 pipeline, offer comprehensive analytical capabilities, provide step-by-step visualization of results, and share a consistent graphical user interface (GUI).

AQuA2’s GUI achieves a notable advancement in user-friendliness compared to its predecessor. We’ve incorporated intuitive functions for imaging analysis that were previously unavailable in AQuA. For instance, AQuA2’s GUI now offers convenient access to capabilities such as image registration, photobleaching correction, and visualization of average curves for manually drawn regions. These additions address the preprocessing needs and result examination that were absent in the AQuA GUI. Moreover, we’ve enhanced the user-controlled parameters to ensure ease of use and alignment with biological principles. Complex parameters and technical intricacies have been abstracted from the GUI, leading to a smoother user experience overall.

### AQuA2 improves the accuracy and efficiency of AQuA

To verify the improvements in accuracy, we compare the detection outcomes of AQuA and AQuA2 on three datasets. We first take an *ex vivo* calcium recording, which was used in the AQuA paper ^14^, as an example (Figure 2A). The results demonstrate that both AQuA2 and AQuA excel in detecting the strong signal near the soma of the astrocyte while exhibiting similar performance in other regions. This suggests that AQuA2 can be considered a viable alternative for applications currently employing AQuA. We then compare the results on a synthetic dataset with known ground truth (Figure 2B) and a real dataset of astrocyte calcium signals in the spinal cord (Figure 2C). Figure 2B and Figure 2C suggest that AQuA’s performance is suboptimal under low SNR or in the presence of large signals. Notably, in the data presented in Figure 2B, we generated a total of 110 real signals. Its low SNR would render signals challenging to detect correctly. However, AQuA detected over 800 events in this scenario, 88% are false positives or negatives, while the error rate for AQuA2 is 7%. Comparisons based on the three datasets suggest that AQuA tends to produce a considerable number of false positives or inaccurately split real signals into trivial fragments while also omitting some significant signals. In contrast, empowered by the new top-down framework, AQuA2 demonstrates robust and accurate performance that aligns more closely with human perception even when distinct signals are connected either spatially or temporally.

**Figure 2.**
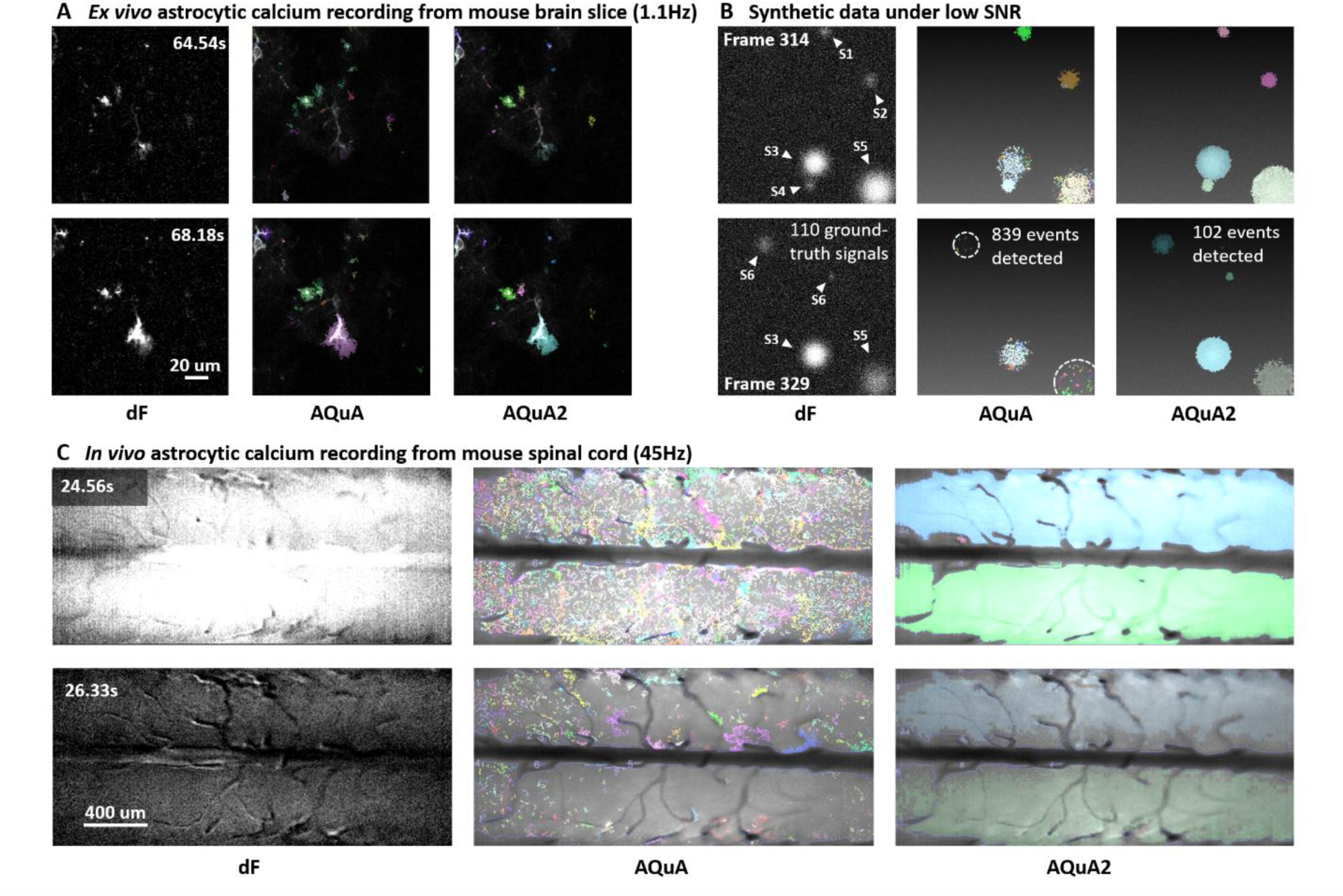
AQuA2 improves the accuracy of AQuA. (A) Performance comparison between AQuA and AQuA2 on a two-photon *ex vivo* astrocyte calcium recording from a mouse slice (recorded at 1.1 Hz), which was used as a test example in the AQuA paper. Similar detection results were obtained, suggesting AQuA2 can be considered a viable alternative for applications currently employing AQuA. (B) Performance comparison on a synthetic dataset comprising large and small signals under a significantly low SNR. Among the 110 ground-truth signals, AQuA2 detects 102 and AQuA detects 105. AQuA2 generates 0 false positives, while AQuA produces 734 false positives. White triangles label the ground-truth events in the present time points. Erroneous detections are marked by white dashed circles. (C) Performance comparison on a one-photon astrocyte calcium recording from the mouse spinal cord (recorded at 45 Hz). During detection, a mask is applied to mitigate the influence of blood vessels. In the AQuA and AQuA2 detection results, each colored region presents one detected signal event. The big events were falsely split by AQuA into numerous fragments. See also supplementary videos.

To demonstrate the efficiency of AQuA2, we compare the execution times across various datasets as illustrated in Table 1. It is worth noting that our comparison focuses solely on the runtime of the event detection pipeline since it occupies over 90% of the overall computation and the runtime of feature extraction may differ based on the demand of users. As illustrated, AQuA2 is typically around twice as efficient as its predecessor. Notably, when signals in the dataset exhibit more complex patterns, AQuA2 demonstrates greater efficiency up to 40-fold. This improvement in efficiency is attributed to two key factors. Firstly, we have adopted the innovative fast algorithm BILCO for the alignment of propagating signals. Secondly, the improved accuracy of AQuA2 reduces the number of false positives, thereby decreasing unnecessary computational time.

**Table 1.** Running time comparison between AQuA2 and AQuA.

| Data name | Data size (pixels x pixels x frames) | AQuA2 | AQuA | AQuA2/AQuA |
| --- | --- | --- | --- | --- |
| ExVivoSuppRaw (Figure 2A) | 502 x 502 x 281 | 114.23s | 355.14s | 32.2% |
| Synthetic data (Figure 2B) | 512 x 512 x 281 | 77.80s | 316.68s | 24.6% |
| Mouse astrocyte calcium (Figure 2C) | 422 x 956 x 350 | 608.84s | 3577.43s | 17.0% |
| Simulated data (case 1)<sup>a</sup> | 512 x 512 x 250 | 125.60s | 198.47s | 63.3% |
| Simulated data (case 2)<sup>a</sup> | 512 x 512 x 250 | 125.17s | 203.89s | 61.4% |
| Simulated data (case 3)<sup>a</sup> | 512 x 512 x 250 | 149.61s | 275.07s | 54.4% |
| GlusnfrSuppRaw<sup>14</sup> | 199 x 108 x 266 | 7.32s | 19.98s | 36.6% |
| Mouse *GRAB-ATP* (Figure 6C) | 256 x 256 x 3372 | 105.32s | 206.35s | 51.0% |
| zebrafish myelin sheath (Figure 6B) | 2028 x 248 x 300 | 84.63s | 3200.10s | 2.6% |

<sup>a</sup> Simulated data used in next section

### AQuA2 outperforms peer methods on simulated data

To further substantiate the accuracy of AQuA2, we carried out a comparative assessment of AQuA2’s performance against AQuA and three other peer tools: suite2p ^12^, CaImAn ^11^, and Begonia ^16^. CaImAn and suite2p are widely used ROI-based methods, while AQuA, Begonia, and AQuA2 are event-based methods. We evaluated these methods using synthetic data under three scenarios frequently observed in real signals: size variability, location variability, and propagation. The performance of each method was evaluated by comparing the detection results with known ground truths. All the methods were fine-tuned to ensure optimal performance. More details about simulation are given in STAR Methods.

Across various scenarios, aside from the specific characteristic we aim to compare, the synthetic data was generated without variation in other features. For example, in the scenario with size variability, there is no location variation and signal propagation. Within each scenario, we initially assessed performance under 10dB SNR and with different degrees of variation - termed ‘size-change odds’, ‘location-change odds’, and ‘propagation frame’, respectively. Then, we evaluate the performance with a moderate variation degree under different SNRs to ensure a comprehensive evaluation. To evaluate the performance, we used two distinct measures: (1) the F1 score, which represents both precision and recall of detections; and (2) the weighted intersection over union (wIoU), which measures the quality of each detection at the pixel level. The details about the two measures are introduced in STAR Methods.

As illustrated in Figure 3, the three event-based methods, AQuA2, AQuA, and Begonia, consistently demonstrate superior performance compared to the ROI-based methods CaImAn and suite2p. This is because ROI-based techniques were initially designed for handling spatially stationary data, making them ill-suited for signals characterized by varying footprints or motion. Among the event-based methods, AQuA2 performs best, consistently achieving F1 scores and wIoU higher than 90% across all experiments and conditions. This highlights that AQuA2 not only exhibits high precision and recall, but also ensures that each detection closely matches its corresponding ground truth, making it an excellent choice for analyzing non-spatially-stationary activities.

**Figure 3.**
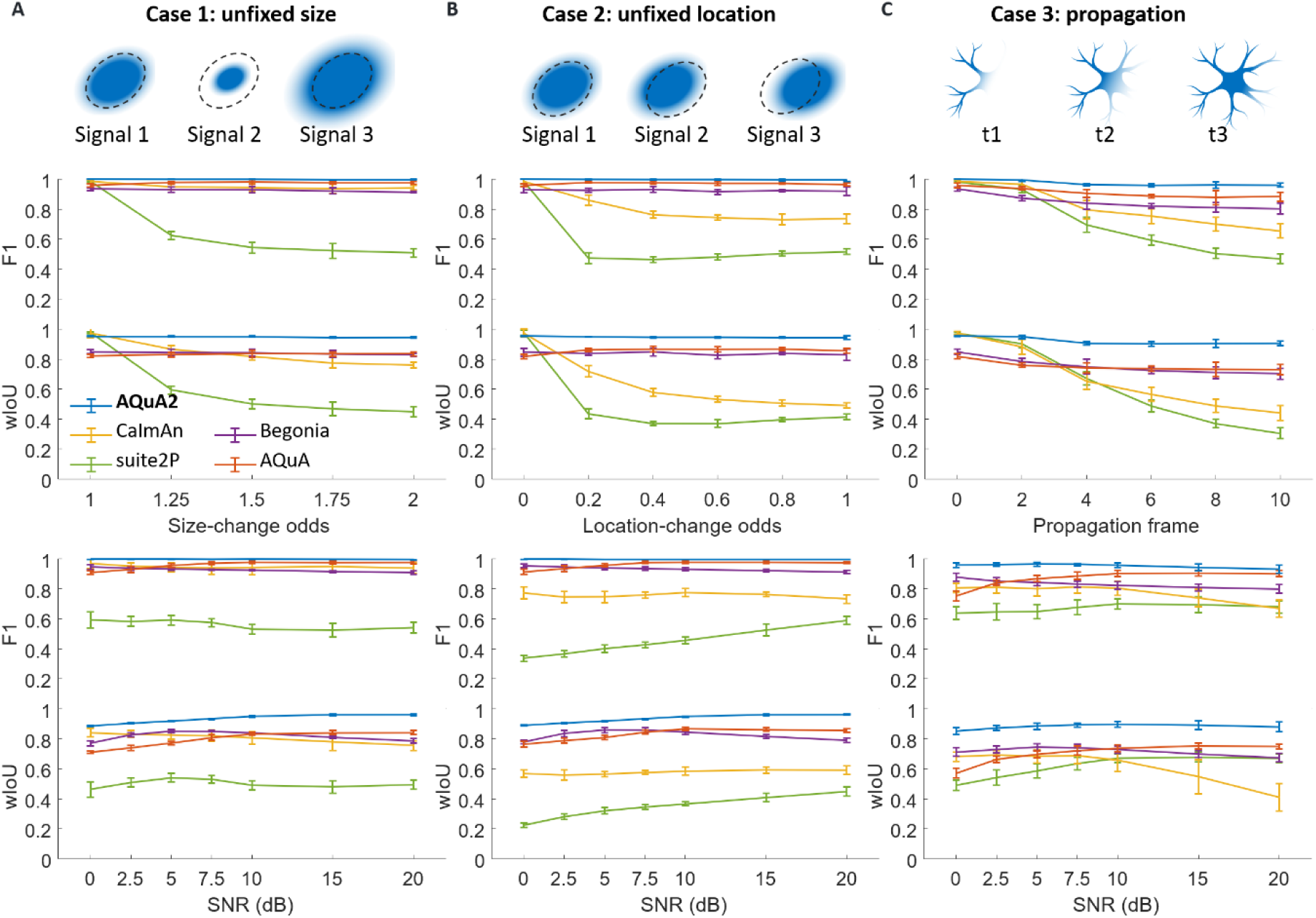
AQuA2 outperforms peer methods (suite2p, CaImAn, AQuA, and Begonia) on simulated data. (A)-(C) Performance (F1 score and wIoU measure, see STAR Methods) comparison between AQuA2 and peer methods under scenarios of unfixed size, unfixed location, and propagation. Top row: Illustration of signal variations. Middle row: Performance comparison under different levels of signal variation under 10dB. Bottom row: Performance comparison under different SNRs with a moderate signal variation. For all results, we used mean ± 2 x standard deviation, derived from 12 independent replications of evaluation. See also Figure S4.

While AQuA and Begonia are better than ROI-based methods, they face practical challenges in real-world applications. The discussion on AQuA has already been provided in the previous section and will not be repeated here. As for Begonia, Figure S5 presents a comparison between it and AQuA2, revealing its significant shortcomings. Begonia is prone to overmerging and artifacts, especially in scenarios with dense signals and low SNR (Figure S5A-B). This is primarily due to its over-simplified assumptions and sub-par modeling, such as splitting connected signals and distinguishing true signals from noises. It is worth noting that we compared functions that were commonly available in peer methods. AQuA2 provides many additional functions that existing peer methods do not offer. For instance, Begonia lacks comprehensive quantification of events, such as the signal propagation analysis (Figure S5C). Automated analysis of functional units and interaction analysis are also lacking in peer methods (Figure S5D).

### AQuA2 identifies signal pattern changes in zebrafish astroglia and neurons under the addition of caffeine

Caffeine, functioning as a central nervous system (CNS) stimulant and an adenosine receptor blocker/antagonist, affects the functions of both neurons and glial cells ^29–31^ and regulates the brain’s internal environment to a certain extent ^29,32^. Recent advancements in imaging techniques allow simultaneous observation of glial and neuronal signals, offering an opportunity to explore caffeine’s impact on the activity of these cell types. However, due to the intricate characteristics of glial cell excitation and the distinct dynamics of neuronal and glial signals, there hasn’t been a universal method for analysis until AQuA2 filled the gap.

Here, we applied AQuA2 to quantify activities and identify functional units of glia and neurons in zebrafish whole-brain. The fish expressed green and red calcium sensors in neurons and radial astrocytes, respectively (*Tg(ELAVL3: GCaMP7f; GFAP: jRGECO1B)).* Zebrafish were engaged in fictive swimming in a virtual-reality (VR) environment with realistic visual feedback during swimming ^33^. It underwent two distinct states: normal and drug (where caffeine was added), as shown in Figure 4A. No additional stimulus was introduced.

**Figure 4.**
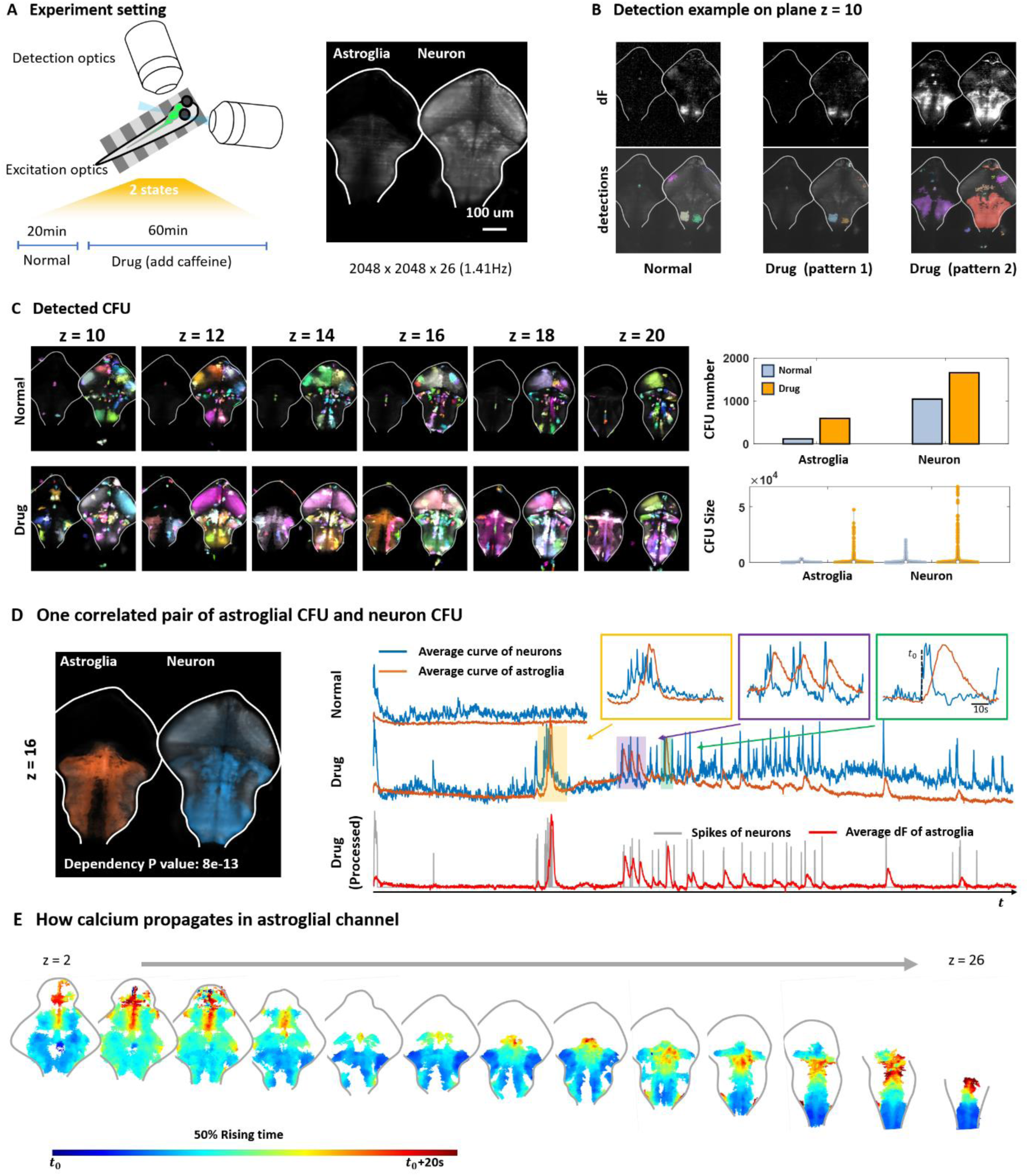
AQuA2 identifies the signal pattern changes in zebrafish astroglia and neurons under the addition of caffeine and reveals the correlation between these two cell types. (A) The experiment setting and average projection of the data. Zebrafish were engaged in fictive swimming in a virtual-reality (VR) environment with realistic visual feedback during swimming, recorded by a light sheet microscope. Experiments of two distinct states were conducted: normal state and under the influence of caffeine (drug state). Astroglia calcium (left) and neuronal calcium (right) were expressed using *Tg(ELAVL3: GCaMP7f; GFAP: jRGECO1B)*. (B) Visualization of dF and AQuA2 detection results under different states. Each colored region represents one detected event. Two distinct signal patterns are found in the drug state. (C) Left: Detected CFUs across different planes under two states. Each colored region represents one extracted CFU. Right: Comparison of number and size among identified CFUs in the two states. (D) The most correlated pair of astroglial CFU and neuronal CFU on one plane, with their curves drawn on the right. Zoom-in traces of three time windows (yellow, purple, green) are also given. For the curves of drug state, neuronal spikes (gray) are additionally extracted (by suppressing small fluctuations on dF) and compared with the average dF of astroglia (red). (E) Propagation pattern of the astroglial calcium signal on different planes. The blue color shows the earliest rising time, while the red color shows the latest rising time.

In Figure 4B, the dF and detection results display representative signals in two states. Notably, in the drug state, there are two distinct signal patterns. Pattern 1 mirrors the normal state, exhibiting solely neuronal signals. This pattern may reflect signal activation due to swimming behavior. In contrast, pattern 2 in the drug state reveals astrocytic calcium signals, often accompanied by a larger neuronal signal. This distinct brain activity pattern indicates a change in brain-wide dynamics resulting from the addition of caffeine.

To conduct a comparative analysis between the two states, we utilized AQuA2 to illustrate the CFUs underlying the signal events across different depths, as illustrated in Figure 4C. The CFUs identified within the hindbrain region are believed to be linked to the neuronal circuits involved in zebrafish swimming, as detailed in Chen et al. ^34^. However, after caffeine addition, a distinct CFU map emerged. While similar CFUs to the normal state were identified, prominent astroglial CFUs and large neuronal CFUs were observed. These findings uncover potential signaling pattern sources activated during brain modulation that may mediate heightened alertness.

### AQuA2 reveals the correlation between neurons and astroglia in zebrafish following the addition of caffeine

Given the pivotal roles glial cells play in supporting, nourishing, and regulating neuronal functions, there has been a growing fascination with the intricate interactions between glial cells and neurons ^35–37^. Emerging evidence suggests that glial cells substantially collaborate with neurons in regulating animal behaviors ^33^. In the previous section, under the influence of caffeine, there were alterations observed in the signaling patterns of both neurons and astroglial cells. Analyzing the functional units of these two cell types can provide a glimpse into the relationship between these two cell types.

To explore the interactions between astroglial and neuronal signals, we utilized AQuA2 to assess the dependencies between astroglial CFUs and neuronal CFUs in the drug-induced state. In Figure 4D, the most notable correlation pair between astroglial CFU and neuronal CFU is depicted alongside their average curves in two states. The average curve of astroglial CFU remains almost constant in the normal state. However, during caffeine treatment, we observed heightened neuronal activity, increased astroglial activity, and an augmented correlation between neuronal and astroglial signals. This correlation is further highlighted when comparing the extracted spikes with the astroglial dF curve, potentially driven by NE after caffeine addition.

To investigate the activation pattern of the astroglial signal, we employed AQuA2 to visualize the propagation patterns of a single astroglial signal across multiple planes, showcased in Figure 4E. Our analysis revealed an earlier activation of signals in specific brain areas like the caudal and lateral medulla compared to those in the more medial regions, which experienced a delayed rising time. These findings shed light on the sources of astroglial calcium activity during caffeine exposure, offering potential insights into the pathways facilitating collaboration between astroglia and neurons.

### AQuA2 unveils sensorimotor signal propagation patterns in the mouse spinal cord

The spinal cord fulfills critical functions in transmitting sensory information, orchestrating movement, and triggering reflex actions ^38–40^. Within the spinal cord, individual laminae contain unique neuronal populations, exhibit distinct connectivity patterns, and serve specialized functional roles ^41,42^. A recently introduced translaminar imaging approach offers the capability for rapid measurements across spinal laminae, providing an opportunity to explore and comprehend the interactions and dynamics inherent to these distinct laminae ^8^. Yet, the absence of a mature analysis tool for this novel imaging technique requires researchers to manually partition the field of view (FOV) into equally sized ROIs for analysis. Given that AQuA2 is an event-based method with versatile applicability, it offers promising and fresh insight into the analysis of such novel data.

We employed AQuA2 to analyze translaminar astrocyte calcium activity in the spinal dorsal horn of *Tg(GFAP:GCaMP6f)* mice. The experimental setup, depicted in Figure 5A, involved securing the mouse on a spherical treadmill and optically recording GCaMP6f activity through a microprism implanted near the lateral edge of the spinal gray matter ^8^. During the recordings, a pressure stimulus was applied to the mouse’s proximal tail. The mouse’s motor behavior was tracked concomitantly (STAR Methods). As illustrated in Figure 5A-C, the pinch elicited noticeable calcium activity within the dorsal regions of the field of view, revealing this spinal region’s response to the sensory stimulus. Within the same recording, AQuA2 also detected two motor behavior-evoked calcium signals that were missed by the ROI analysis mentioned above.

**Figure 5.**
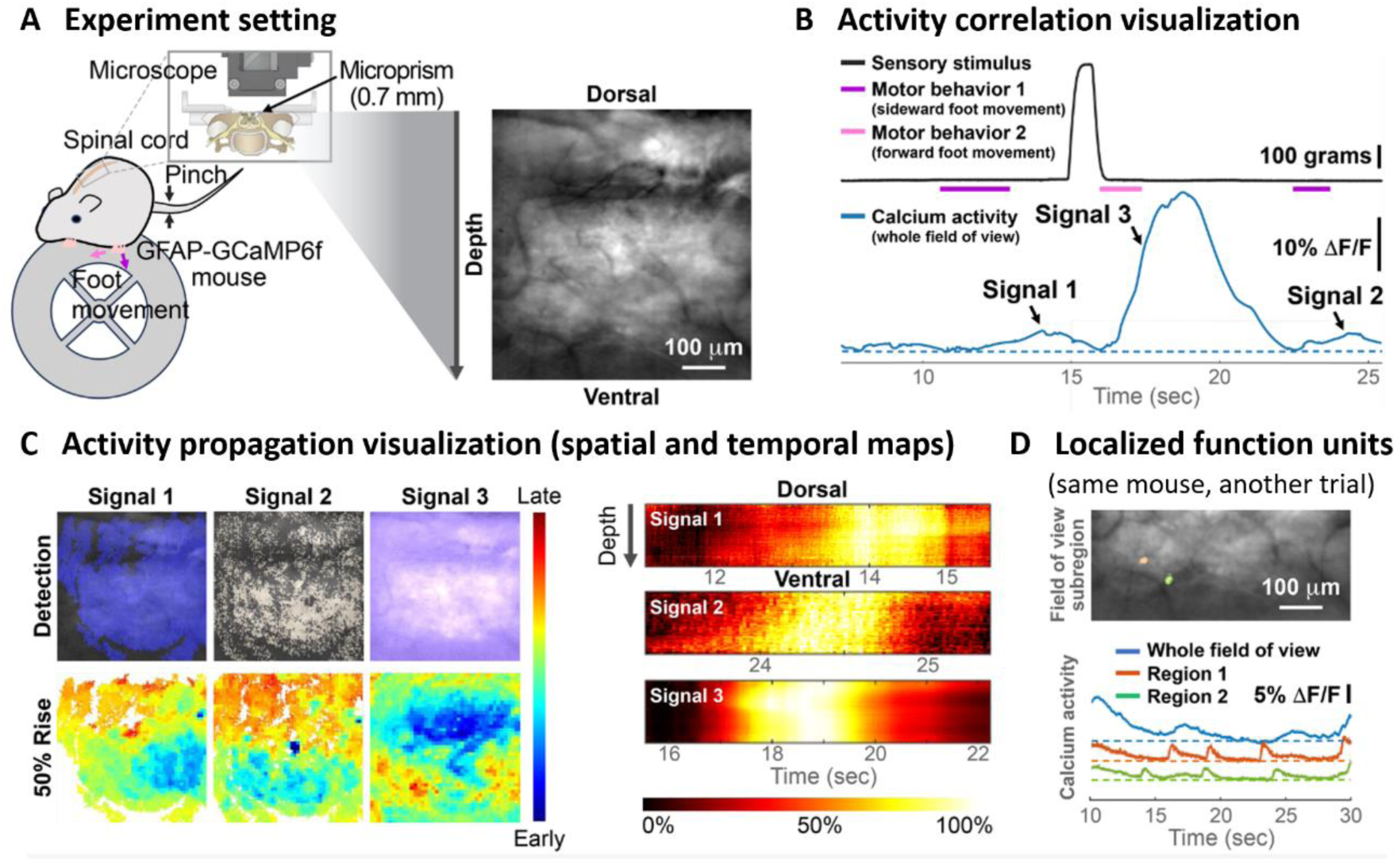
AQuA2 unveils differences in signal propagation of sensory and motor-evoked astrocyte calcium signals in the mouse spinal cord. (A) Experimental setting: Translaminar imaging ^8^ was performed in the lumbar dorsal horn of an awake behaving *Tg(GFAP:GCaMP6f)* mouse on a spherical treadmill. A pressure stimulus was applied to the mouse’s proximal tail. (B) Temporal relationship between the applied sensory stimulus (tail pinch; black), mouse motor behavior (sideward and forward foot movements; purple and pink, respectively), and the average calcium activity across the field of view (blue). (C) Spatial (left) and temporal (right) activity propagation maps of the three signals indicated in B. The color spectrum depicts earlier activation in blue and later rising times in red. (D) Detected localized functional units (orange and green; top) and their corresponding calcium transients (bottom).

To determine the distinguishing features between these sensory and motor-evoked signals, we utilized AQuA2 to visualize their respective signal propagation patterns. As depicted in Figure 5C, the sensory-evoked signal displayed a propagation pattern starting from the dorsal side, while the motor-evoked signals emanated from the ventral side. The findings suggest that peripheral sensory stimuli and motor actions engage neural circuits in distinct spinal laminae, that astrocytes’ activity patterns reflect this neural activity, and that the astrocyte excitation can propagate across spinal laminae on a seconds timescale in behaving mice. This discovery has implications for how astrocytes might regulate spinal neural circuit activity or plasticity and underscores AQuA2’s ability to uncover new biological phenomena leading to testable hypotheses about cell or circuit function.

Additionally, beyond the translaminar signals, Figure 5D illustrates the detection of local signals within spinal laminae of the same mouse during a different trial. These signals, often missed by other methods, imply the existence of localized functional units, akin to those observed in the brain.

### AQuA2 quantifies signals across biosensors, cell types, organs, animal models, and imaging modalities

The recent advancements in indicators and imaging techniques have facilitated the recording of diverse neurotransmitter and neuromodulator activities including ATP ^6^, glutamate ^43^, GABA ^44^, serotonin ^45^, NE ^5^, and others, significantly aiding research into brain mechanisms. Despite this progress, signals of numerous molecular sensors lack dedicated analysis methods. AQuA2 will serve as a universally applicable tool for filling this gap.

To validate the ability of AQuA2 to quantify and analyze signals of various types, we conducted tests on datasets encompassing a variety of animals, cell types, molecular sensors, organs, and imaging modalities. Due to space constraints, only four applications are depicted in Figure 6. These datasets include microglia calcium recordings in the spinal cord of a mouse, oligodendrocyte calcium imaging in zebrafish, recordings of ATP signals in the mouse visual cortex, and dual-color recordings of NE signals and glial calcium in the whole brain of zebrafish.

**Figure 6.**
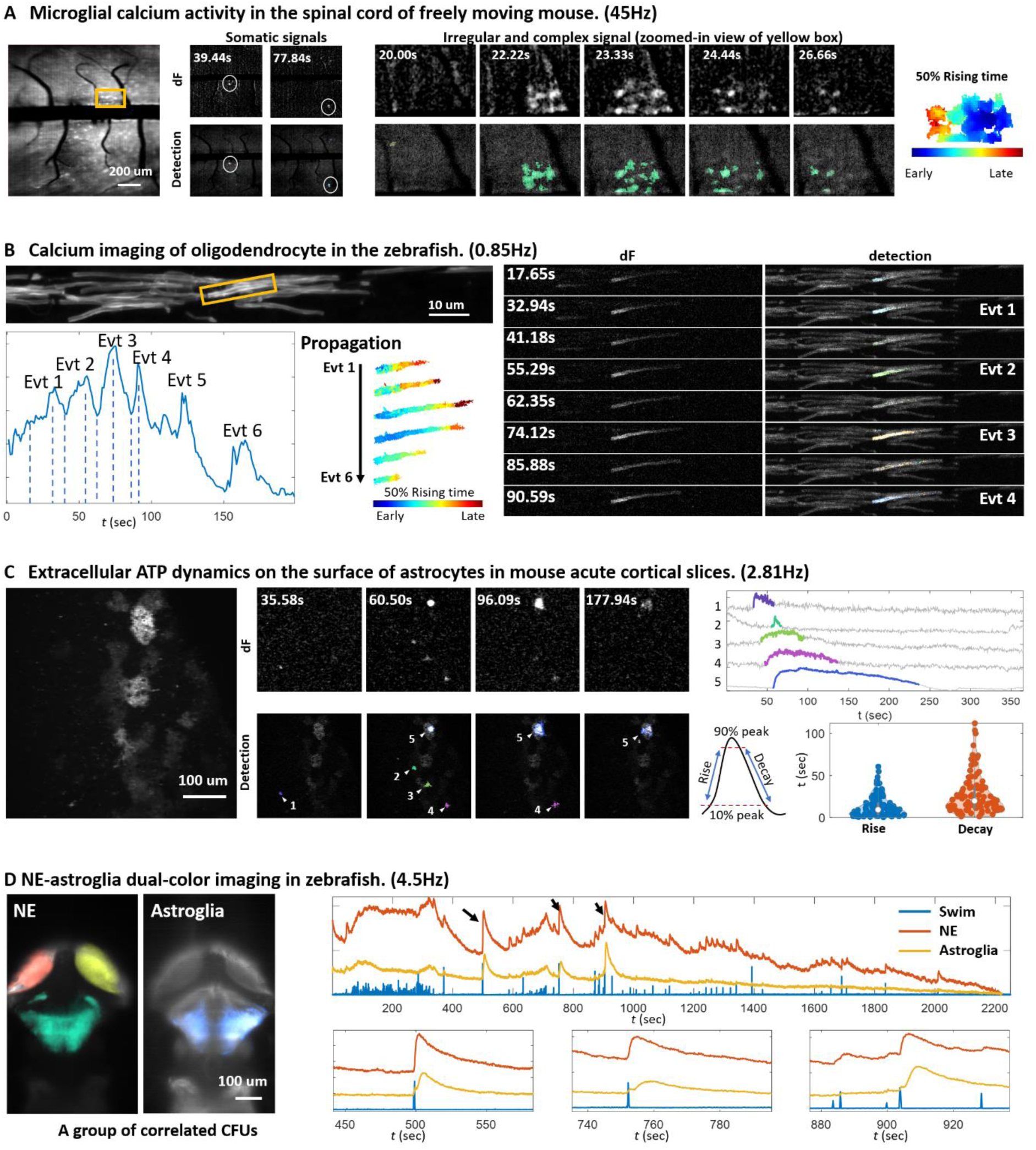
AQuA2 quantifies signals across biosensors, cell types, organs, animal models, and imaging modalities. (A) Application of AQuA2 to quantifying the signals from mouse spinal microglia (expressed by *Tg(CX3CR1:GCaMP5g)*) imaging data. The comparison between dF and detection for both circular signals and the spatially complex signal (shown in a zoomed-in view of the selected orange box) is provided. On the right, the quantified propagation pattern of the spatially complex signal is given, with blue denoting early rising time and red denoting late rising time. (B) Application of AQuA2 for accurate quantification of distinct oligodendrocyte calcium dynamics within the CNS myelin sheath of zebrafish. The calcium was expressed through *Tg(mbp:memGCaMP7s)*. The average curve of the region of interest (labeled by yellow) is depicted in the bottom-left. In the bottom middle, the propagation pattern of events is showcased, with earlier activation represented in blue and later rising times in red. On the right, a comparison between dF and AQuA2 results is provided, with each colored region representing a distinct detection. (C) Application of AQuA2 for the quantification of extracellular ATP dynamics, captured by the *GRAB-ATP* sensor ^6^ and two-photon microscope, on the surface of astrocytes in acute cortical slices. Example representative events (marked by white triangles) are presented with their corresponding average curves (event duration is indicated by the event color). Statistical analysis of event temporal features (Rising duration and decay duration) is provided on the bottom-right. (D) Application of AQuA2 for identifying the swim-related regions on a light sheet norepinephrine (NE)-astroglia dual-color recording of zebrafish. NE was expressed by *Tg(ELAVL3: GRABNE)* and astroglial calcium was expressed by *Tg(GFAP: jRGECO)*. On the left, each colored region represents a swim-related CFU. On the right, swim strength (blue) and average curves of two channels (red and yellow) are depicted. Three zoom-in figures are provided for three signals. See STAR Methods for experimental details.

AQuA2 can detect microglial calcium activity, as shown in Figure 6A. These transients were recorded in the superficial dorsal horn of behaving *Tg(CX3CR1:GCaMP5g-tdTomato)* mice using a wearable macroscope ^46^. Both somatic signals and signals with irregular shapes and complex spatiotemporal features were identified. AQuA2, therefore, can detect signals with distinct spatiotemporal characteristics in the same dataset, including localized and propagating cellular activity.

AQuA2 was also employed to detect and visualize temporally dense oligodendrocyte calcium activities within CNS myelin sheaths. The calcium sensor was expressed around the membrane using the zebrafish line *Tg(mbp:memGCaMP7s).* As illustrated in Figure 6B, the calcium transients distinctly exhibit signal propagation along the myelin sheath and form interconnected patterns in the temporal dimension. The propagation and interconnectedness pose a challenge for other analysis tools. However, by leveraging AQuA2, each transient was effectively isolated and quantified, accompanied by a clear visualization of its propagation pattern. This successful application underscores the capability of AQuA2 to handle intricate and complex signals.

We also tested how AQuA2 detects biological signals other than calcium. Using AQuA2, we quantified extracellular ATP dynamics captured by the *GRAB-ATP* ^6^ sensor on the surface of astrocytes in acute cortical slices. As depicted in Figure 6C, ATP signals exhibit heterogeneous temporal characteristics, ranging from long-lasting (hundreds of seconds) to extremely brief (less than ten seconds). Despite this heterogeneity, AQuA2 successfully detects them using the same parameter settings, showcasing its ability to detect signals with diverse temporal characteristics.

The relationship between NE signals and astroglial calcium signals within the brain is also an area of investigation for biological researchers. Employing AQuA2 on a dual-color dataset in zebrafish, which captures NE and astroglial calcium signals in separate channels (*Tg(ELAVL3: GRABNE; GFAP: jRGECO)*) and records zebrafish swim commands via electrophysiology, we identified and visualized swim-related CFUs, as depicted in Figure 6D. Upon analyzing the curves across different channels, the results demonstrate a notable occurrence: following zebrafish swim spikes, NE and astroglial signals manifested in pairs. This observation is consistent with behavior-generating circuits engaging both NE signaling and astroglial calcium activity.

In addition to the mentioned applications, further use cases are shown in Figures S6 and S7, encompassing zebrafish acetylcholine signals, zebrafish dopamine signals, and zebrafish dual-color data featuring ATP and neuronal calcium signals, among others. Collectively, AQuA2 can be effectively utilized to quantify and analyze signals across a variety of molecular sensors, cell types, organs, animal models, and imaging modalities, consistently providing highly informative results.

## DISCUSSION

While advanced genetic fluorescent probes and imaging techniques have unleashed the potential for neurotransmitter, neuromodulator, and other molecular-level studies, the corresponding analysis tools have not progressed at the same pace. This situation necessitates researchers devising their own analysis pipelines or attempting to adapt mismatched analysis tools. In response to this need, we developed AQuA2, a completely new version of AQuA with improved accuracy, efficiency, versatility, and many new functions.

Compared to the existing approaches, AQuA2 can quantify and analyze a more diverse range of signals and low SNR scenarios, a capability that AQuA lacks. For instance, as illustrated in Figure 2B and Figure 2C, the limitations of AQuA’s framework become evident when the target signal occupies a larger spatial area, leading to the frequent misidentification of broad signals as multiple unclear small events - an issue mentioned by Rupprecht et al. ^47^. Additionally, AQuA also struggles to effectively detect signals with longer durations, while AQuA2 successfully identifies signals >100s (Figure 6C). AQuA may erroneously identify any short temporal fluctuations as events, even when the duration of the longer target signal is presumed, particularly in low SNR scenarios (Figure 2C). AQuA2 addresses these issues by embracing a novel framework, incorporating prior knowledge in the early stage, and making a set of technical innovations. As a result, AQuA2 can perform flexible detection across diverse biosensors, cell types, organs, animal models, and imaging modalities, verified by our real applications, as shown in Figure 6 and Figure S7.

The development of the CFU analysis effectively bridges the gap between ROI-based and event-based methods. It serves as a more flexible version of ROI, tolerating the variation between signals activated from the same source. This flexibility is crucial because, in many datasets, the territories of signals emanating from the same source are often not fixed. That will violate the assumption of ROI-based methods, while event-based methods lack the ability to analyze the region behind events. The CFU module fills this gap, enabling the analysis of functional units behind events with variation. To probe the functional relationship between CFUs, a new statistical approach was developed, taking event occurrence sequences as input. This allows AQuA2 to reveal interrelationships between functional units with different waveforms, such as neuronal CFU with spike signals and astroglial CFU with bell-shaped signals (Figure 4D). Interactions between CFU and stimulus/behavior can also be analyzed (Figure 6D), and the estimated dependency could serve as a distance measure to group CFUs into potential circuits (Figure S7D).

Propagation analysis, a key feature of AQuA, is seamlessly integrated and optimized in AQuA2. It employs joint alignment techniques to match all time points among pixel curves and the reference curve. In contrast to the correlation-based approach, which relies on matching the entire curve and is consequently vulnerable to waveform variations, leading to suboptimal results, joint alignment analysis offers greater flexibility and the ability to provide rich, precise information, as shown in Figure S2A. However, the long computation time and large memory usage have been obstacles in using AQuA for many users. This challenge is particularly pronounced when dealing with spatially large events, where an increased number of pixels escalates the problem’s scale.

For instance, in the past, propagation analysis like Figure 5C would lead to memory overflow on our workstation as the event occupies the whole field of view. The adoption of our new algorithm, BILCO ^24^, has solved this problem. Compared with the propagation analysis in AQuA, it now completes the analysis in only 1/10th of the time and 1/10th of memory usage, enabling AQuA2 to efficiently examine the propagation patterns of spatially large signals.

In addition to the improvements discussed above, AQuA2 enhances accessibility and usability. Three versions of the same pipeline were implemented for different users, each offering unique advantages. For instance, the MATLAB version allows direct code modification and debugging, enabling researchers to adjust the analysis for their data. The Fiji plugin version supports integration with other Fiji plugins and usage across different systems (Windows, Linux, and Mac) without requiring environment installation. AQuA2-Cloud offers online analysis, offloading computations to a remote server. Furthermore, AQuA2 adopts clear and user-friendly parameters, eliminating previously confusing settings in AQuA and ensuring ease of use without significant usability barriers.

We have demonstrated numerous applications of AQuA2 across various challenges, encompassing the quantification of diverse signals, visualization of signal propagation patterns, and investigation of interactions among different signal types in dual-color data. However, the capabilities of AQuA2 extend beyond these tasks. For instance, with the advancing technology, 3D fluorescent data will likely offer more insights and become the next trend. Although our paper primarily presents analyses on 2D images, AQuA2 has also been validated on 3D inputs, as depicted in Figure S3. These applications highlight AQuA2 as versatile software for comprehensively analyzing diverse fluorescent imaging data, facilitating a broad spectrum of scientific inquiries.

### Limitations of the study

While we anticipate widespread usage of AQuA2, it is important to acknowledge its limitations. The detection pipeline predominantly relies on discerning the temporal patterns of signals, demanding a stable fluorescence baseline. Although the pipeline can accommodate gradual changes in the baseline, significant jitter or distortion in the image may inevitably lead to false positives. This is commonly observed when the animal suddenly moves but can be mitigated using image registration techniques. For stereotypical neuronal nuclear or somatic calcium signals, AQuA2’s performance may not exceed that of ROI-based methods, as ROI assumptions align well with neuronal dynamics. But for neuropil analysis, AQuA2 may perform better, as the assumptions of ROI-based methods may not hold, given the possible compartmental signals and the near-impossibility of achieving perfect stabilization at the axon/dendrite-scale tissue level. AQuA2 assumes that signals are positive deviations from the baseline. Theoretically, signals with negative deviation from the baseline can also be modeled with AQuA2 by inverting them. However, further studies are warranted if a biosensor reports positive and negative signals simultaneously.

## STAR METHODS

### KEY RESOURCES TABLE

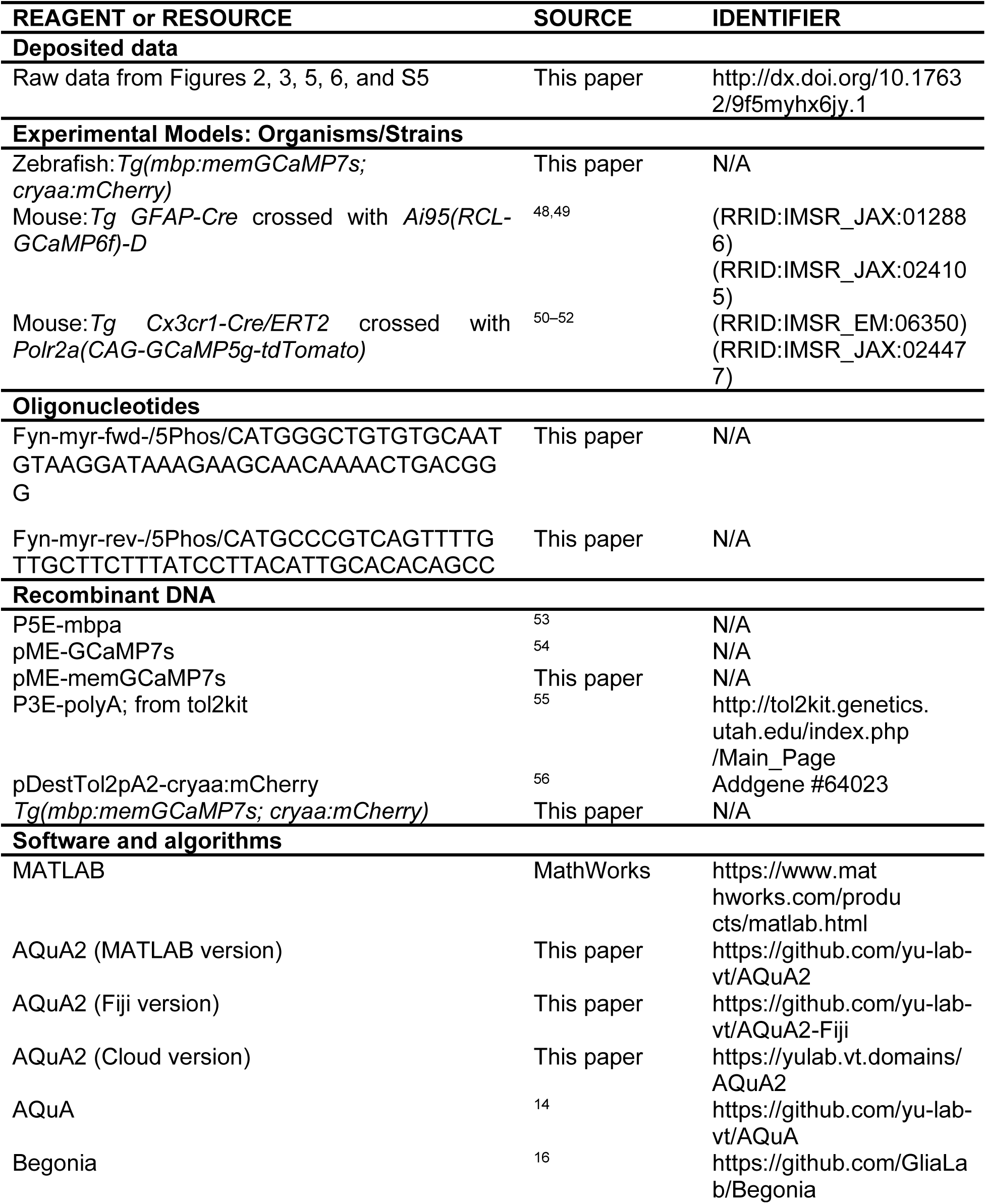
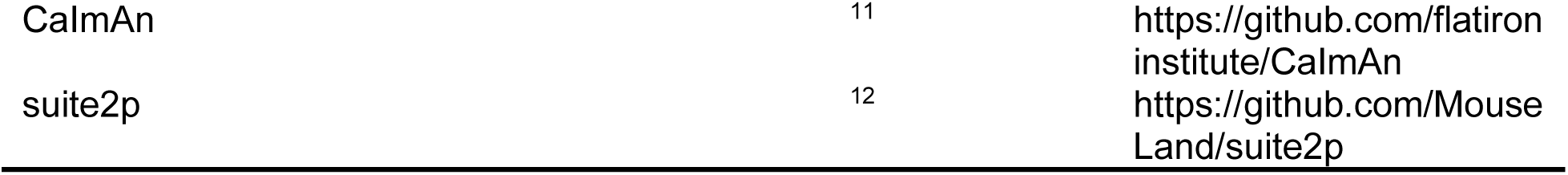

### RESOURCE AVAILABILITY

#### Lead contact

Further information and requests for resources and reagents should be directed to and will be fulfilled by the lead contact, Guoqiang Yu.

#### Materials availability

Zebrafish lines generated in this study will be available upon request from Misha Ahrens or David Lyons.

#### Data and code availability

- The raw data reported in this paper are either shared through Mendeley with the dataset identifier http://dx.doi.org/10.17632/9f5myhx6jy.1 or will be shared by the lead contact upon request due to the size limit of Mendeley.
- All original code has been deposited at https://github.com/yu-lab-vt/AQuA2.
- Any additional information required to reanalyze the data reported in this paper is available from the lead contact upon request.

### EXPERIMENTAL MODEL AND SUBJECT DETAILS

#### Animals

##### Zebrafish (Ahrens lab)

Experiments were conducted according to the guidelines of the National Institutes of Health and were approved by the Standing Committee on the Use of Animals in Research at Harvard University. Animals were handled according to IACUC protocols 22-0216 (Ahrens lab). We did not determine the sex of the fish we used since it is indeterminate at this age. Fish were raised in shallow Petri dishes and fed ad libitum with paramecia after 4 dpf. Fish were raised on a 14 h:10 h light:dark cycle at around 27°C. All experiments were done during daylight hours (4–14 h after lights on). All protocols and procedures were approved by the Janelia Institutional Animal Care and Use Committee.

##### Zebrafish (Lyons lab)

Adult zebrafish were housed by the bioresearch and veterinary service at the Queens Medical Research Institute at the University of Edinburgh. Studies were carried out with the approval of the UK home office and according to its regulations (project license PP5258250). Adult animals were maintained at a 14-h day and 10-h night cycle. Embryos were housed at 28.5°C in 10 mM HEPES buffered E3 medium. Zebrafish embryos were imaged at 3 dpf, before the onset of sexual differentiation. In this study the transgenic (Tg) *mbp:memGCaMP7s* was used.

##### Mouse (Nimmerjahn lab)

All mouse procedures followed the National Institutes of Health (NIH) guidelines and were approved by the Institutional Animal Care and Use Committee (IACUC) at the Salk Institute under protocol number 13-00022. Mouse strains used in this study included GFAP-Cre (RRID: IMSR_JAX:012886), Ai95(RCL-GCaMP6f)-D (RRID: IMSR_JAX:024105), CX3CR1-Cre/ERT2 (RRID: IMSR_EM:06350), and Polr2a(CAG-GCaMP5g-tdTomato) (RRID: IMSR_JAX:024477) mice. All mice were on a C57BL/6 J background. Mice were group-housed, provided with bedding and nesting material, maintained on a 12-h light-dark cycle in a temperature (22±1°C) and humidity-controlled (45–65%) environment, and provided with food and water ad libitum. The experiments involved 6- to 12-week-old heterozygous male and female GFAP-GCaMP6f mice (Figures 2C, 5) and 15- to 16-week-old male and female CX3CR1-GCaMP5g-tdTomato mice (Figure 6A). Cre-mediated DNA recombination in Cre/ERT2 mice was induced following established protocols. Briefly, tamoxifen (T5648, Sigma-Aldrich) was diluted in corn oil (C8267, Sigma-Aldrich) and injected intraperitoneally at 100 mg per kg (body weight) once a day for five consecutive days, followed by imaging ∼3–5 weeks after the final injection. Sex as a biological variable was not considered in the research design and analyses, as the study’s primary goal was to demonstrate the analysis approach’s technical capabilities. Experimental mice used in individual experiments typically originated from different litters. Mice had marks for unique identification. No criteria were applied to allocate mice to experimental groups.

##### Mouse (Poskanzer lab)

All experiments were done in *Mus musculus* with a Swiss or C57BL/6J genetic background. The mouse protocols were approved by the University of California, San Francisco Institutional Animal Care and Use Committee (IACUC). Mice were housed on a 12:12 light-dark cycle and were provided food and water ad libitum. Approximately equal numbers of male and female mice were used in experiments.

#### Transgenesis

##### Generation of the Tg(mbp:memGCaMP7s; cryaa:mCherry) transgenic line

To generate a *Tg(mbp:memGCaMP7s; cryaa:mCherry)* transgenesis construct, we first created a tol2kit-compatible middle-entry vector containing the coding sequence for the membrane-tethered memGCaMP7s, pME-memGCaMP7s. To do this, we digested our previously generated plasmid pME-jGCaMP7s ^54^ (containing an untethered, cytoplasmic GCaMP7s coding sequence) at the start codon with NcoI-HF enzyme (GCCA<u>CC**ATG**G</u>, NcoI recognition sequence underlined, start codon in bold, NcoI-HF from New England Biolabs). Into this digested vector we then ligated two annealed primers, Fyn-myr-fwd and Fyn-myr-rev, phosphorylated at the 5’ end (from IDT DNA Technologies), which encode the myristoylation motif of human Fyn kinase flanked by overhanging NcoI-compatible ends. The sequence for this plasmid was verified by Sanger sequencing.

To generate the final Tol2 expression construct *Tg(mbp:memGCaMP7s; cryaa:mCherry)*, we then recombined 10fmol of the following entry vectors: previously described 5’-entry vector 5E-mbp ^53^ ME-memGCaMP7s, and 3E-polyA from the tol2kit ^55^; and 20fmol of destination vector pDestTol2pA2-cryaa:mCherry ^56^ (Addgene #64023), using LR Clonase II Plus. 3-4 clones were tested for correct recombination by restriction enzyme digestion.

To establish a stable transgenic line, we injected 5pg of *Tg(mbp:memGCaMP7s; cryaa:mCherry)* plasmid DNA with 50pg tol2 transposase mRNA into wild-type zebrafish eggs at the one-cell stage. This yielded memGCaMP7s-expressing oligodendrocytes in injected embryos, which were then raised to adulthood. Founders were identified by screening their F1 offspring for germline transmission using the cryaa:mCherry marker.

##### *GRAB-ATP* virus injections

Viral injections were done in P0-3 pups. For all injections, pups were anesthetized on ice for 3 minutes and then positioned on a digital stereotax. Using a microinjection pump (UMP-3, World Precision Instruments), a mixture of *AAV9-GfaABC1D-GrabATP*1.0 with dye (2*μ*l virus + 0.5*μ*l Fast Green dye; virus titer 1.78× 10^13^ VG/mL) was injected at four injection sites in a 2×2 grid over the visual cortex. The first injection coordinate was 0.8-0.9mm lateral and 1.5-1.6mm rostral from lambda, and each site was separated by 0.8mm. At each site, 20-30nl of the virus was injected at two z-depths (0.1 and 0.2mm below the skull) at 3nl/s. Pups recovered from anesthesia on a heating pad before being returned to their home cage.

### METHOD DETAILS

#### Data collection

##### Mouse *ex vivo* GCaMP imaging using two-photon microscopy

The *ex vivo* GCaMP dataset used in Figure 2 is a public dataset recording the astrocytic calcium signals of the mouse brain slice. Its description can be found in the AQuA paper ^14^ and it can be downloaded from https://drive.google.com/file/d/13tNSFQ1BFV__42TY0lZbHd1VYTRfNyfD/view.

##### Zebrafish whole-brain imaging with a light sheet microscope (Ahrens lab)

Zebrafish whole-brain recordings were captured using a custom light sheet microscope with adjustable z resolution and consistent x and y resolutions of 0.40625um/px, as described by Mu et al. ^33^. Dual-color recordings involved simultaneous scanning at wavelengths of 488 nm and 561 nm, resolution of 2048 x 2048 pixels, with images separated using a dichroic mirror and captured on an Orca Flash 4.0 v2 camera. The detection objective was a 16x/0.8NA lens.

During experiments, paralyzed zebrafish performed fictive swimming in a VR environment, with the drifting gratings providing visual feedback, as shown in Figure 4. For the experiment in Figure 4, green and red calcium signals were expressed in neurons and astroglia using *Tg(ELAVL3: GCaMP7f)* and *Tg(GFAP: jRGECO1b)*, respectively. Caffeine 1mM was added in drug state. The frame rate was 1.41 Hz, with 10um per plane resolution, and 26 planes were collected. For the experiment in Figure 6D, the green NE signal and red astroglial calcium signal were expressed using *Tg(ELAVL3: GRABNE)* and *Tg(GFAP: jRGECO)*, respectively, in an open-loop setting with no visual feedback provided regardless of swim commands collected. The frame rate was set at 4.5 Hz. One suction electrode (∼60 um inner diameter) filled with external solution, was placed over the dorsal side of the fish’s tail and attached with gentle negative pressure. The voltage signal recorded by this tail was amplified and filtered (band-pass 300 Hz - 3 kHz) with a MultiClamp 700B amplifier. This signal was then smoothed through convolution with an exponential filter and used as the ‘swim signal’.

##### Mouse spinal recordings (Nimmerjahn lab)

Mouse spinal cord imaging was performed as previously described. Briefly, animals were implanted with a spinal and head plate under general anesthesia approximately one week before laminectomy and received Buprenorphine ER/SR (0.5 mg/kg) to minimize post-operative pain.

For intralaminar imaging through a dorsal glass window (Figures 2C, 6A), a laminectomy (2 mm wide × 4 mm long) was performed at the T12-T13 vertebra level, corresponding to spinal segments L3–L5. The dura mater overlying the spinal cord was kept intact, and a custom-cut #0 coverslip was used to seal the laminectomy, creating an optical window for imaging.

For translaminar imaging through an implanted glass reflective microprism (Figure 5), a 0.7 mm × 0.7 mm × 0.7 mm microprism with aluminum-coated hypotenuse (cat. no. 4531-0021; Tower Optical) was UV-cured (NOA 81; cat. no. 8106; Norland Products Inc.) to a custom-cut #0 coverslip matching the intended laminectomy size (3 mm wide × 4 mm long). A laminectomy (3 mm wide × 4 mm long) was performed at the T13-L1 vertebra level, corresponding to spinal segments L4–L6. Using a dissecting knife (cat. no. 10055-12; Fine Science Tools) attached to a stereotactic arm, a small incision was made 0.7 mm lateral to the central vein’s center, coinciding roughly with the interface between the dorsal root ganglia (DRG) and spinal white matter in adult mice. The incision extended 0.7 mm in the rostrocaudal direction and 0.7 mm in depth, matching the microprism dimensions. No spinal cord tissue was removed. No blood exuded from the incision site upon retraction of the dissecting knife. The microprism implant was positioned above the incision site and slowly lowered until fully inserted (∼0.7 mm depth). Excess fluid was removed using sterile absorbent paper points (cat. no. 50-930-669; Thermo Fisher Scientific). The implant was affixed to the surrounding bone with instant adhesive (cat. no. 3EHP2; Grainger).

Fluorescence image data were acquired with a wearable multi-color microscope (Figure 5) or macroscope (Figure 2C, 6A) using 473 nm excitation by a fiber-coupled DPSS laser. For imaging through the dorsal optical window (Figures 2C, 6A), using the wearable macroscope on freely behaving mice (16” x 16” open field), the typical average light power at the tissue surface was <250 μW mm^−2^. For imaging through the implanted microprism (Figure 5), using the multi-color microscope on focally restrained mice, 275-325 μW mm^−2^ was used. Translaminar image data were acquired four weeks after microprism implantation and at around 75 μm focal depths from the vertical microprism-tissue interface, where 0 μm was defined as the point when cells or blood vessels first came into focus. Our recordings showed no signs of phototoxicity at the specified light powers, such as a gradual increase in baseline fluorescence, lasting changes in activity rate, or blebbing of labeled cells. All data were acquired at the image sensor’s full resolution (1280 × 960 pixels) and maximum frame rate (∼45 Hz). The image data were preprocessed as previously described, including illumination correction, background subtraction, and image registration. In a subset of recordings (Figures 2C, 5), mechanical stimuli were delivered to the animal’s tail using a rodent pincher system (cat. no. 2450; IITC Life Science, Inc.). Pinch pressures were applied in the dorsoventral direction approximately 6 mm from the base of the animal’s tail. Synchronously acquired analog data included the pressure sensor output from the rodent pincher system and the on-off TTL signal of the wearable microscope’s light source, which were recorded at 1 kHz using DAQExpress 2.0 software (National Instruments). Pinch application and mouse behavior were also recorded on a video camera (≥20 Hz; Stingray F-033, Allied Vision Technologies) using AVT SmartView software (v1.11). To synchronize imaging with video data, we placed a near-infrared LED within the video camera’s FOV, triggered from the microscope’s light source drive signal (TTL pulse). The video data were cropped to the LED-on period and scored manually regarding mouse motor activity, pinch onset, and offset.

##### Zebrafish oligodendrocyte calcium recording (Lyons lab)

In the experiment depicted in Figure 6B, calcium in oligodendrocytes of zebrafish was expressed around the membrane using the *Tg(mbp:memGCaMP7s)*. The expression was recorded using an LSM880 confocal microscope operating in airy scan fast mode, with a 20x water immersion lens. The zebrafish were then subjected to imaging for a total of 300 frames and 10 stacks. The acquisition frequency was set at 0.85 Hz, and the resolution was 0.0595373um per pixel. Then the data underwent several processing steps, including airy scan processing, maximum intensity projection, and registration using turbo reg. The final processed data had dimensions of 2028 pixels x 248 pixels x 300 frames.

##### Mouse *GRAB-ATP* recording (*ex vivo* acute slices) using a 2P microscope (Poskanzer Lab)

In the experiment depicted in Figure 6C, acute coronal slices from the visual cortex (300um thick) were collected from P28-42 mice, as described previously by Pittolo et al. ^57^. All imaging was done at room temperature on a custom 2P microscope using a 920 nm laser, 525/50 emission filter, 256×256 pixel resolution, 20x objective with 1.0 N.A. (Olympus), a spatial resolution of 2.08um/px, and a frame rate of 5.62 Hz. Before 2P imaging, each slice was incubated for ∼5 minutes in a 20mL recirculating standard artificial cerebrospinal fluid (ACSF) bath with 1uM TTX +/- 500nM POM1. Then, *GRAB-ATP* dynamics were recorded for 5 ∼ 6 minutes. Before AQuA2 detection, all videos were smoothed along the z-axis by averaging every 2 frames.

##### Synthetic data generation

For the synthetic data utilized in contrasting the performance of AQuA and AQuA2, we employed randomly generated signals, each exhibiting varying SNR. For the synthetic data used to compare all the peer methods, we simulated datasets based on templates derived from real signals. To emulate authentic brain activity, we employ three distinct synthetic settings: unfixed size, unfixed location, and propagation. These datasets were generated under a consistent 10dB SNR. Consequently, across these scenarios, we evaluate performance under varying SNRs: 0dB, 2.5dB, 5dB, 7.5dB, 10dB, 15dB, and 20dB—resulting in a total of 6 distinct settings. Within each setting, we generate 4 diverse ground truth patterns and incorporate random noise for each pattern repeated 3 times.

##### Definition of SNR

We define the SNR of simulation using the following formula:

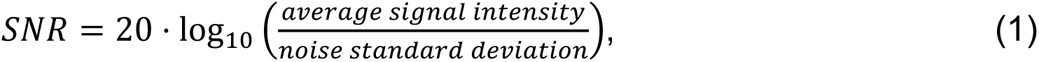

where the average signal intensity is the average intensity of all pixels belonging to the ground truth.

##### Synthetic data for comparing performance of AQuA and AQuA2

We simulated the synthetic data used in Figure 2B with size 500 pixels x 500 pixels x 500 frames, and a background image that changes gradually in the y dimension. We initially generated 100 signal centers at random positions, then applied Gaussian filters with uniformly distributed sizes on both spatial and temporal dimensions to them. As a result, those signals have radii ranging from 10 to 40 pixels, durations ranging from 10 to 20 time points, and centers with intensity ranging from 10% to 20% dF/F0. Afterward, we generated 10 signals using a similar approach in the same dataset, this time incorporating large Gaussian filters to simulate scenarios involving both large and small signals. The large signals have radii ranging from 40 to 80 pixels, durations ranging from 80 to 150 frames, and centers with intensity ranging from 30% to 40% dF/F0. Large signals are allowed to overlap with small signals, thus some small signals may be obscured. Finally, Gaussian noise was added, with the variance equal to the baseline value on each pixel to model the phenomenon of photon collection through microscopy. A rough estimation of the SNR across the whole dataset is around −10dB.

##### Synthetic data for comparing performance of Begonia and AQuA2

We simulated the synthetic data used in Figure S5B with size 500 pixels x 500 pixels x 300 frames, and a background image that changes gradually in the y dimension. We generated 300 signal centers at random positions, then applied Gaussian filters with uniformly distributed sizes on both spatial and temporal dimensions to them. As a result, those signals have radii ranging from 10 to 40 pixels, durations ranging from 10 to 20 time points, and centers with intensity ranging from 10% to 20% dF/F0. Gaussian noise was added, with the variance equal to twice of the baseline value on each pixel. A rough estimation of the SNR across the whole dataset is around −16dB.

##### Spatial template of the signals for peer method comparison

Given that ROI-based methods often demand users to define signal sizes, we utilized the regions of neuronal signals in mice as spatial templates, since the sizes of these signals are relatively consistent. These templates underwent processing involving morphological closing and dilation to fill gaps and refine boundaries. The templates are then randomly translated and rotated on a mask of size 512 pixels x 512 pixels, with the constraint that the closest distance between two ROIs should be larger than 5 pixels. On the final mask, there are 66 spatial templates, each with a size close to 20 pixels x 20 pixels.

##### Temporal template of the signals for peer method comparison

Similarly, ROI-based methods typically require the specification of the decay constant, which implies a fixed temporal pattern. To enable these methods to achieve relatively good performance, we adopted a signal template with a fast intensity increase and slow decay. The template reaches its maximum intensity of 1 at the third time point, with the first two frames reaching 0.4 and 0.8, respectively. Then the template presents a decay following the function *e*^−0.3*t*^, where t is the time point after the peak. In general, if we consider a threshold of 20%, the temporal duration of the template is approximately 8 time points.

##### Ground truth data generation for peer method comparison

We generated ground truth data of size 512 pixels x 512 pixels x 250 frames, using both spatial templates and temporal templates. For each spatial template, we uniformly distributed the peaks of the temporal pattern with a density of 0.04. To prevent signals from being too close for distinction, we retained only the first signal if two peaks were located within a time window of length 5 frames. Furthermore, to ensure that each signal is fully contained within the data, we do not allow the peaks to be in the initial or final frames. Following convolution with the temporal pattern at peak positions, each spatial template generates approximately 7-8 signal events. Furthermore, within the synthetic settings featuring variable sizes, diverse locations, and propagation, to guarantee the distinguishability of each ground-truth event, we prohibit signal centers from overlapping. The center, in this context, is defined as the portion exhibiting an intensity surpassing 50%.

For ground-truth labeling, a watershed algorithm based on the signal centers is used, with the constraint that pixels labeled as ground truth must have an intensity greater than threshold 0.2. In general, each synthetic data contains approximately 400-500 events.

##### Synthetic setting - unfixed size for peer method comparison

To simulate potential changes in the size of signal events, we randomly altered the size of signals that correspond to the same spatial template while keeping their spatial center unchanged. A parameter called “size-change odds” controls the maximum size change, where a “size-change odds” of x means that the size can randomly vary from 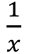 to x times its original size. Dilation and shrinkage have equal chances of occurring. The parameter varies from 1 to 2 in increments of 0.25. To achieve this, we generated a uniformly random variable between 1 and the parameter to represent the scale factor and another uniformly random variable between 0 and 1 to determine whether to dilate or shrink, with a threshold of 0.5. When testing the robustness under different SNRs, we set the “size-change odds” to be 1.5.

##### Synthetic setting - unfixed location for peer method comparison

To simulate potential changes in the location of signal events, we randomly moved the position of signals that correspond to the same spatial template to a nearby region. The “location-change odds” parameter controls the maximum distance of translation, where an odds of x means that each signal event may be shifted within a range of 0 to x times the template diameter. The angle of translation is also random. In our setting, the parameter varies from 0 to 1 in increments of 0.2. To achieve this, we generated two uniformly random variables: one ranging from 0 to the parameter to represent the translation distance, and another ranging from 0 to 2*π* to represent the rotation angle. When testing the robustness under different SNRs, we set the “location-change odds” to be 0.4.

##### Synthetic setting - propagation for peer method comparison

In our simulation of signal propagation, we generated signals with two distinct types of potential propagation: “move” and “grow”, with equal chances of occurring. For the “move” propagation type, the signal patterns of pixels remain the same, but with a gradually changing delay among adjacent pixels. As a result, the signal appears to move to another position. Similarly, in the “grow” setting, there is also a gradually changing delay, but the signal patterns of pixels within the same event differ in duration. Pixels that produce signals earlier have a longer duration achieved by interpolation, and all pixels within the same event end at the same time point. Thus, such signals appear to grow into a larger region.

For both propagation types, we set the propagation speed to be the same, at 0.15 times the radius of the spatial template. This means that the patterns of pixels with a distance of 0.15 times the radius will be delayed by one time point. The propagation direction is fixed within one event but is selected uniformly from 0 to 2*π*. Additionally, all signals have the same propagation length, denoted by the “propagation frame” parameter in our setting. In our simulation, the parameter varies from 0 to 10 in increments of 2. When testing the robustness under different SNRs, we set the “propagation frame” to 4.

##### Synthetic data for comparing correlation-based method and joint alignment for propagation analysis

We generated synthetic data with dimensions of 500 pixels x 500 pixels × 100 frames, featuring a row-dependent waveform and column-dependent delay for each pixel. Before the peak, the waveform follows a Gaussian shape for each pixel, while post-peak, it exhibits an exponential decay with a parameter linked to the row. Typically, the duration of the curve ranges from 30 frames (at the top row) to 70 frames (at the bottom row). Concurrently, the curves of pixels experience varying delays across different columns, ranging from 0 (at the leftmost) to 20 (at the rightmost). Subsequently, noise was introduced to achieve an SNR of 0 dB in the synthetic data. During the application of the correlation-based method, we utilized the average curve of the entire dataset as the reference curve.

##### AQuA2 event detection pipeline

To enhance the accuracy, efficiency, and versatility of AQuA, we have re-engineered the event detection pipeline using advanced machine-learning techniques. The pipeline comprises five steps as shown in Figure S1A, with the first four being essential for detecting signals, the fifth being optional for detecting temporally global signals, and the final step being used to quantify the detections. Step 1 (Preprocessing) addresses experimental artifacts such as motion and photobleaching effects, removes baseline fluorescence, and estimates noise variance. After this step, *dF*/*σ* is displayed, where dF represents dynamic fluorescence and *σ* denotes the noise standard deviation. Step 2 (Active region detection) applies a statistical test to identify active regions, each of which potentially contains activity. As each active region may encompass multiple signals, Steps 3 (Temporal segmentation) and 4 (Spatial segmentation) are designed to segment these regions so that each result adheres to our signal event assumption of containing only one peak and originating from a single source. Step 5 (Global signal detection) is optional and involves detecting global signals based on removing the signal events detected in the first four steps.

##### Step 1: Preprocessing in AQuA2

The preprocessing step in AQuA2 tackles image registration, photobleaching correction, baseline estimation, and noise modeling, aiming to prepare well-processed data for subsequent analysis. Due to the optional nature of image registration and photobleaching correction, we have placed their introduction towards the end.

###### Baseline estimation in preprocessing

The raw fluorescence F consists of three components: baseline *F*_0_, signal *S*, and noise *N*. Baseline refers to the background signal present in the image. It does not reflect changes in molecular concentration, thus lying outside the scope of interest. Therefore, accurate estimation of the baseline is a crucial step to mitigate baseline impact on the analysis of target signals.

Traditional baseline estimation methods often rely on a single projection of the imaging data as the baseline, disregarding potential temporal variations. This oversight can result in false detections. For example, if the baseline steadily decreases over time, using a constant baseline would inevitably lead to overestimation in certain time intervals and underestimation in others. Therefore, it’s imperative to account for the trend of baseline changes for accurate signal detection.

To distinguish between baseline and signals, we presume that the baseline represents the slow-changing component along the temporal dimension, while target signals change faster. This contrast serves as the foundation for baseline estimation. Under the assumption of a slow-changing baseline and non-negative signals, we model the baseline for each pixel by a piece-wise linear function: Initially, we apply a moving average filter (typically with a window of 25 time points) to mitigate the impact of noise. Subsequently, we segment each pixel’s time course into intervals (the default segment length is 200 time points). By linking the minimum point of each time segment, we derive a piecewise linear function representing the trend. To counterbalance the bias introduced by selecting the minimum point, a predetermined quantity is then added.

###### Noise modeling in preprocessing

Once the baseline is isolated, the preprocessing step focuses on modeling the noise to effectively identify the target signals. While many methods typically estimate a single variable to represent the noise standard deviation for the entire dataset ^12,14^, this may not accurately capture the reality. In fluorescent data, the noise level can vary significantly across different locations, with brighter pixels exhibiting stronger noise while the noise in darker regions is comparatively weaker. That’s one reason why researchers usually use *dF*/*F*_0_ to represent signal strength.

Taking this phenomenon into account, we initially estimate the noise at the pixel level. We operate under the assumption that the noise at different time points on the same pixel follows an independent and identically distributed Gaussian distribution. Given that real signals might lead to an overestimation of noise, we estimate noise by calculating the square of the difference between adjacent time points with the following equation rather than directly computing variance.

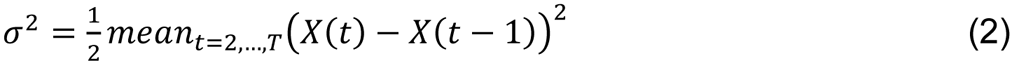

The noise estimation, as described above, may not be entirely accurate due to the limited temporal length of each pixel and an insufficient number of samples. Thus, an estimation leveraging information from all pixels is anticipated to enhance robustness. Given that fluorescence recording conforms to an optical process following a Poisson distribution, there exists a pronounced correlation between the noise variance and the expectation value (baseline) ^58^. We propose harnessing this correlation by employing a linear function to model the relationship between variance and baseline value, thus integrating information from all pixels for enhanced estimation accuracy.

Recognizing that recording may be influenced by truncation and saturation, leading to different relationships at both low and high baseline levels, we employ a three-segment piecewise linear function instead of a simple linear function (Figure S2B). The modeling of this function can be efficiently solved by transforming the determination of the endpoints of each segment into a shortest path problem within a constructed graph (Figure S2C). In this graph, each pixel is treated as a sample/node and organized into several layers, where each node in the graph represents a potential endpoint. For instance, the connection between node *u* in layer 1 and node *v* in layer 2 signifies one segment in the piecewise linear function with these two nodes as endpoints. To ensure the piecewise linear function is unidirectional, we sort the nodes based on their baseline values. Only edges between a node *u* and the next-layer node *v* with a higher baseline value (*F*_0_(*v*) > *F*_0_(*u*)) are permitted. To minimize the fitting error between samples and the piecewise linear function, we assign the weight of an edge as the cumulative absolute error between samples and the piecewise linear function, calculated as 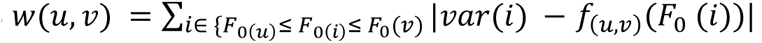. Here, *var*(*i*) and *F*_0_(*i*) represent the variance and baseline value of the *i*_*th*_ node, and *f*_(*u*,*v*)_(*F*_0_(*i*)) is the estimated variance of the baseline value *F*_0_(*i*) on the segment between nodes *u* and *v*. We designate the first and last layers to comprise only a few nodes close to the minimum and maximum baseline values, respectively, enabling automatic determination of the overall start and end points from these nodes.

By solving the shortest path within this constructed graph, which can be achieved with linear complexity, we establish a robust relationship between baseline and noise variance. This integration of baseline information enables us to obtain a more reliable noise estimation, enhancing the accuracy of our analysis.

###### Optional image registration in preprocessing

In fluorescent imaging data, it’s common to encounter sudden movements of animals leading to jitter or drifting in the recorded data. These artifacts often manifest as rigid translations, given the limited field of view in microscopy. To mitigate motion artifacts, AQuA2 provides the option to utilize the cross-correlation algorithm ^59^ to align each frame with a reference image. By default, the reference image is determined by averaging the first 10 frames.

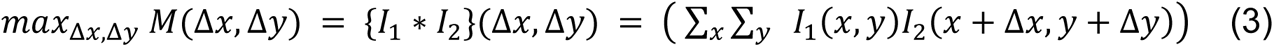

where Δ*x*, Δ*y* represents the translation in two dimensions, *M* shows the similarity, * denotes the convolution symbol, and *I*_1_ and *I*_2_ represent two images.

This algorithm calculates the optimal shift between two images by identifying the maximum similarity among all possible translations, where similarity is indicated by the summation of two images’ dot product. While it may appear necessary to iterate through every element in the image for each (Δ*x*, Δ*y*) to compute similarity, the convolution theorem in the Fourier domain offers a highly efficient solution ^59^. Once the optimal shift is determined, rectifying the translation involves shifting the images in the opposite direction.

###### Optional photobleaching correction in preprocessing

Photobleaching is another common artifact observed in fluorescent imaging data, wherein fluorescent probes gradually degrade due to exposure to excitation light, leading to a loss of fluorescence capability. Such decay effect is typically modeled by an exponential function ^60^. AQuA2 offers two approaches to model photobleaching decay: (1) Global Modeling: Decay is modeled globally on the average curve of all pixels by fitting the decay with the model *f*(*t*) = *ae*^−*bt*^ + *c*. Then, the fluorescence intensity is divided by the decay to remove the photobleaching influence; (2) Intensity-based Correction: Acknowledging that the bleaching rate may vary across pixels with different intensities, an intensity-based correction method is employed. Pixels are clustered based on their baseline intensity, and decay is modeled on the average curve of each cluster.

##### Step 2: Active region detection in AQuA2

With *F*_0_estimated in the preceding step, we can derive dF, which exclusively comprises components, signals denoted as S, and noise labeled as N. Due to the unknown distribution of signal intensity, Bayesian decision-making cannot be employed to determine the optimal threshold for distinguishing between signal and noise. Hence, we leverage the non-negativity of signals and employ hypothesis testing to detect active regions. *dF*/*σ* serves as a z-score, indicating the likelihood of a voxel belonging to a signal or not, where *σ* denotes the noise standard deviation. By applying thresholding based on a user-defined parameter on *dF*/*σ*, we can filter out voxels more likely to be noise. The remaining voxels, exhibiting high z-score, are aggregated as potential active regions.

Notice that people may have some prior knowledge about the target signals according to the experiment setting. The information on the size (default set at 20 pixels), duration (default set at 5 time points), or shape of the signals (default set the circularity constraint at 0) could be introduced as filters. Recognizing that bright pixels are typically more reliable, we utilize a top-down multiple thresholding strategy to incorporate this prior knowledge. At each threshold level, regions above the threshold are examined using the filters. Candidate regions that pass the filter criteria are retained as active regions (regions with lower intensity will override those with higher intensity).

##### Step 3: Temporal segmentation in AQuA2

Recall that an event is defined as a spatiotemporally connected region characterized by fluorescent dynamics, exhibiting a single peak pattern and originating from a single source. In the preceding step, spatiotemporally connected regions potentially containing target signals are detected. However, within a single active region, multiple signal events may appear. Therefore, segmentation becomes necessary.

In this step, AQuA2 performs temporal segmentation on the active regions to ensure the presence of a single peak pattern. This is achieved through the following procedures: identifying regions with significant peak patterns as seeds, expanding these seeds into subregions, and merging subregions with similar temporal patterns. The resulting regions after temporal segmentation are termed super events, each exhibiting only one peak pattern. For convenience and without loss of generality, in this step, we work on the score map which is dF normalized by the estimated noise standard deviation.

###### Identifying regions with significant peek patterns as seeds

To ensure that each result contains a single peak pattern, the active region is segmented by first identifying potential peak patterns. This is accomplished by assessing the temporal significance of candidate regions selected using a top-down, multi-threshold, and multi-scale strategy.

The candidate region selection begins with a top-down multi-threshold strategy. This approach is chosen because regions under a low threshold may contain multiple peak patterns, similar to the active region, whereas candidate regions above a high threshold are more likely to contain only one peak pattern.

The multi-scale strategy employed for region selection involves considering large-scale yet weak signal regions, achieved through downsampling operations. This aids in aggregating information over a broader field of view while enhancing SNR. Additionally, this strategy incorporates spatial morphology considerations. For instance, if two candidate regions possess identical size and intensity but differ in spatial morphology—one resembling a complex winding line and the other having a regular shape like a square or circle—intuitively, the latter should hold greater significance. By aggregating information from neighboring regions, this strategy integrates morphology considerations. By default, AQuA2 considers scales of 2×2, 4×4, and 8×8.

To ensure the singularity of the result’s peak, if any seeds have already been detected within the selected candidate region, AQuA2 will preserve these seeds without reassessment. Otherwise, the temporal significance of the candidate region is evaluated. Only if the significance is sufficiently high will the region be deemed to contain a peak pattern and thus be utilized as a seed.

To measure the temporal significance of a candidate region, AQuA2 avoids direct evaluation on its average curve. This is because potential signal propagation and the presence of other signals could disrupt the significance assessment, making it difficult to establish a suitable time window for evaluation. Instead, AQuA2 measures the temporal significance at each pixel and combines the pixels’ significances to evaluate the region’s significance. The significance of each pixel is assessed based on the contrast between the selected time window and neighboring time points. Considering the nature of peaks, the contrast significance is evaluated twice: once comparing the selected window with the preceding neighbors, and another time comparing it with the subsequent neighbors.

For each pixel, assuming the intensities of the selected time window are (*x*_1_, *x*_2_, …, *x*_*n*_), and the intensities on the time window with a certain length before and after are (*x*_*n*+1_, *x*_*n*+2_, …, *x*_*n*+*m*_), where n is the number of foreground pixels and m is the number of background pixels (by default, n=m), the contrast *L* can be calculated as follows:

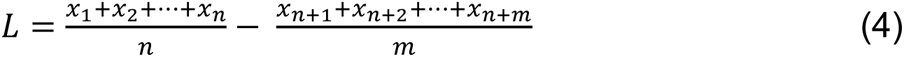

Given that the selection of candidate regions is biased by the thresholding operation, AQuA2 employs order statistic theory to evaluate temporal significance. This approach assumes that both foreground and background are sampled from the same null hypothesis distribution, and thus the bias and variance of the contrast L can be calculated given the order information.

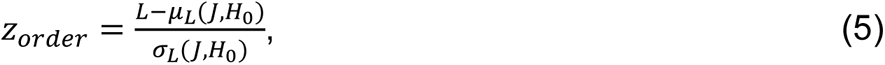

where J is the relative order of foreground group and background samples, *H*_0_ represents the null hypothesis distribution (here we assume it is Gaussian distribution), *μ*_*L*_(*J*, *H*_0_)and *σ*_*L*_(*J*, *H*_0_) denote the calculated bias and standard deviation under order J and null hypothesis *H*_0_.

After determining the significance of each pixel’s curve, the next step involves aggregating these values to evaluate the significance of the entire region. However, since each pixel corresponds to a different duration, it’s crucial to assign varying weights to their significance. Pixels with longer durations typically provide more reliable data and therefore warrant greater weighting. Thus, we can calculate the region’s significance:

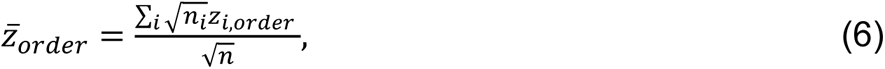

where *z̅*_*order*_ is the region’s significance and *n*_*i*_ is the length of the selected time window of ith pixel. Such a combination strategy assigns a duration-based weight *n*_*i*_ for each pixel. It is consistent with treating each intensity sample within ith pixel as having a score *z*_*i*,*order*_ such that the significance of the sample group for *i*th pixel is 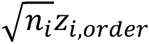.

We have backgrounds on two sides for each pixel, and thus finally two significant scores for each selected region. To pass the seed significance test, both the scores of two sides should be larger than a user-defined threshold (the default value is 3.5 due to multiple testing).

After completing the seed detection step, it’s plausible that no seeds are found in certain active regions. This absence suggests that these regions lack sufficient temporal significance and may represent false positives. We will consequently eliminate these regions. Hence, the seed detection step doubles as a significance filter, aiding in the removal of temporally non-significant active regions. By eliminating such regions early in the process, we enhance the reliability of subsequent analyses and ensure that only genuinely significant regions are further considered.

###### Growing seeds into subregions

If one active region contains multiple seeds, we expect to segment the region into “subregions”. Each subregion should be a grown region from a seed. Although multiple region growing methods could be applied, we adopt a marker-controlled watershed algorithm due to its high efficiency and consistency with intuition. To implement this marker-controlled watershed, we first reverse the *dF*/*σ* and then set the values of all seeds to be negative infinity. The part outside the active region is also treated as another seed, and the boundary values of the active region are set to be positive infinity. As the watershed algorithm runs on this processed map, nearly all the pixels within the active region will be assigned to some seed except the final ridges between adjacent subregions. To eliminate these ridges, we conduct an additional round of subregion growth to fill the gaps.

###### Merging subregions with similar temporal patterns

Considering the potential over-segmentation of the active region due to the texture or gap within the fluorescent image, AQuA2 employs a merging step to merge the neighboring subregions with a similar peak pattern. The obtained results are termed “super events”.

Two rules for merging subregions are used: spatially neighboring subregions with similar signal patterns should be merged (the similarity is introduced in the following), while temporally adjacent subregions with large spatial overlap (measured by intersection over union, IoU) should not. Based on these rules, within one active region, AQuA2 greedily searches for neighboring subregion pairs with the smallest temporal dissimilarity and merges them if their spatial overlap is tolerable until the temporal dissimilarity exceeds a certain threshold.

Since subregions are grown solely based on intensity, unexpected intensity fluctuations among *dF*/*σ* values can lead to irregular shapes. Consequently, certain subregions may contain small portions of frames with pixels outside the primary region. Using the footprint of the region may therefore overestimate spatial overlap, potentially preventing the merging of subregions that should be combined. Moreover, after region growing, some subregions may persist longer than the majority time window, allowing additional peaks to become involved. This complicates the comparison of signal patterns. To address this issue, we extract representative or majority subregions to ensure that the most significant and prevalent features of each subregion are accurately captured. These representative or majority parts will replace the subregions during the merging process.

To measure the temporal dissimilarity of two patterns x and y, dynamic time warping (DTW) is used to find the best matches for each time point. Given the obtained optimal warping function P={(*k*_*x*,*i*_, *k*_*y*,*i*_)}, where i is the index of the path, *k*_*x*,*i*_ and *k*_*y*,*i*_ represent the corresponding time points on x and y respectively, the average distance of each pair of corresponding time points can be calculated:

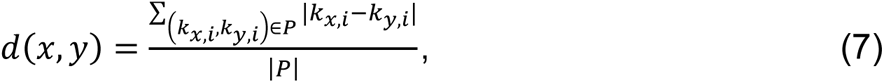

where |P| represents the size of the path P.

However, *d*(*x*, *y*) is an absolute distance measure, which can be easily affected by the signal duration and is not suitable to serve as a dissimilarity measure. Thus, AQuA2 normalizes the distance by the minimum duration among x and y, then uses the following metric as the dissimilarity measure between signal patterns. This final distance metric incorporates information on delay, shape difference, and duration.

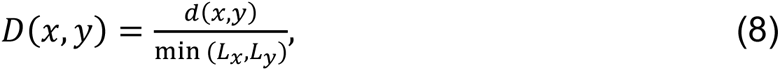

where *L*_*x*_ and *L*_*y*_ denote the lengths of time windows for signal patterns x and y.

##### Step 4: Spatial segmentation in AQuA2

Although a super event is supposed to be a spatiotemporal connected region with a single peak pattern, the definition of the event also requires the event to originate from a single source. We assume that in an event, the signal source is the location where the signal is first activated. Therefore, the signal source can be detected by propagating information. The spatial segmentation in AQuA2 consists of two major procedures: propagation estimation and the extraction and expansion of signal sources.

###### Propagation estimation

To determine the potential propagation or delay at each pixel, it is intuitive to identify the reference shift that maximizes correlation with the pixel’s curve. However, despite similarities in patterns, gradual deformation within the super event can render propagation information unreliable, as depicted in Figure S2A.

Acknowledging the potential for deformation, we utilize alignment techniques to identify the optimal match or warping path between the pixel’s curve and the reference (the average temporal pattern from the largest subregion). Among alignment techniques, AQuA2 employs joint alignment, which integrates structural information from the data and has demonstrated superior performance ^24^. To enhance efficiency, joint alignment is solved using the BILCO algorithm ^24^, developed by our team, which achieves a 10-fold improvement in both time and memory usage.

Before applying the BILCO algorithm, several detailed operations are performed. For all pairs to be aligned, the time course is normalized by its noise standard deviation, and its corresponding reference is scaled to the same range for correct alignment. This takes into account the quality of the time series, assigning a larger weight to those with a high SNR. Additionally, to further improve efficiency, a downsample operation is performed on super events with a large size. For example, if the spatial size of one super event is 4000 pixels, a 2 ×2 downsample operation will be performed.

###### Segmentation based on propagation

AQuA2 employs a multi-threshold strategy, which is executed in ascending order of 50% rising time, derived from propagation information, to identify potential source regions. The contrast between the candidate region and its surroundings is then calculated to determine the source. The system offers ten levels of sensitivity for signal source detection, with level 10 being the most sensitive and level 1 being the least.

There would be three cases in a super event: (1) No signal source region is detected. Then the super event is like a stationary signal activity and there is no need to segment. (2) Only one signal source is found. It denotes the super event comes from the same signal source, and thus still no need to segment. (3) Multiple signal sources are extracted, which means the super event is indeed coming from different sources. Here, we will assign each pixel to a different source using a marker-controlled watershed algorithm.

It’s crucial to emphasize that while our objective is to spatially segment the super event to ensure each final event originates from a single source, this segmentation shouldn’t rely solely on pixel labels. Due to the potential interference in the data, doing so may lead to the center of one region being labeled as one event while its surroundings are considered as another event. To prevent such occurrences, we aggregate the intermediate results from step 3, the subregions, to form the final event based on the majority source label.

##### Step 5: Global signal detection in AQuA2

Global signal detection is an optional step aimed at identifying signals of extended duration that may have been obscured by signals detected in earlier stages. For instance, if a signal component is the sum of 10 temporally disjoint local signals, each with a duration of 10, over a global signal in the same time window with a duration of 100, then the previous steps would only be able to detect signals with shorter durations. The detection of the global signals is achieved by first removing the impact of local detections, where the intensities of the voxels in the local detections are interpolated according to the temporal boundaries of each detection, and then repeating the previous steps with a different duration limitation.

###### AQuA2 CFU identification

With the intuition that regions exhibiting repetitive signals are more likely to bear biological significance, we introduce the concept of a CFU. A CFU is defined as a spatial area generating a set of events with similar locations and morphologies. Events associated with a specific CFU share similar spatial territories, although not necessarily identical. The CFU concept combines the strengths of both ROI-based and event-based methodologies. It retains the notion of a “region” from ROI, grouping events with spatial consistency, while incorporating the flexibility of event-based methods to allow for some spatial variation among signals.

This concept serves as a more generalized version of ROI, offering increased flexibility. Unlike ROI, which is spatially static and confines signals to precisely the same territories, CFU acknowledges that signals may exhibit slight spatial variations. This flexibility is crucial for exploring functional units, as signals often do not maintain exactly the same territories in most data. Signals with higher intensity will span larger territories, while those with lower intensity will occupy relatively smaller areas. ROI-based methods may segment the union territory of these signals into multiple parts, identifying the overlapped region as a core ROI and the surrounding regions as independent ROIs, leading to the identification of false positive ROIs. The flexibility inherent in CFU positions it as a promising solution for exploring the functional units within such datasets.

CFU can be conceptualized as a collection of events characterized by consistent spatial patterns. Intuitively, identifying CFU can be framed as a clustering method. Among classical unsupervised clustering algorithms, AQuA2 adopts a hierarchical clustering approach. This choice is motivated by the advantage of utilizing the interpretable largest allowable inconsistency as a hyperparameter, as opposed to relying on a predetermined cluster number - a challenging aspect to ascertain. In the task of exploring functional units, the unknown number of functional units makes methods like K-means hard to apply, and there is the potential for false positive events to be clustered into distant regions. However, the hierarchical clustering approach ensures that only signals with similar territories are clustered together. Even with potential false positive events, a filter based on the number of signals associated with CFU can effectively eliminate such instances.

Using the hierarchical clustering approach, the keys in CFU identifications are how to define the spatial territory of one event and how to estimate the consistency/inconsistency between two territories. Each event is a set of voxels {*x*_*i*_, *y*_*i*_, *t*_*i*_} and we can reformulate the coordinates into the form {*p*_*j*_, *n*_*j*_}. Here, *i* is the voxel index, *j* is the pixel index, *p*_*j*_ represents the spatial location of one pixel, and *n*_*j*_ denotes the count of voxels located at the same *p*_*j*_ position. Inspired by the fact that a pixel with a longer duration *n*_*j*_ holds greater importance, we calculate a weighted spatial set {*p*_*j*_, *w*(*p*_*j*_)} to demonstrate the spatial territory for each event,

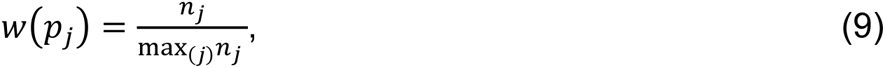

where *w*(*p*_*j*_) is the weight assigned to the pixel *p*_*j*_.

Having obtained the spatial territories of signal events, our subsequent step involves estimating the consistency between any pair of two events. Common measures like cosine similarity or Pearson’s similarity necessitate input vectors of the same dimension. However, our spatial territories may comprise weight parameters with varying sizes. While an option is to fill the weights with zeros for pixels outside the event, this approach has its drawbacks. As each event may only occupy a small fraction of the field of view, the filled weights will dominate the vectors and unreliably inflate the consistency. Here we selected the weighted version of Jaccard similarity ^61^ to handle this task. This choice accommodates the varying sizes of spatial territories and provides a more accurate measure of consistency. Its unweighted version, commonly known as intersection over union, is a standard measure for evaluating the match between two objects in computer vision tasks.

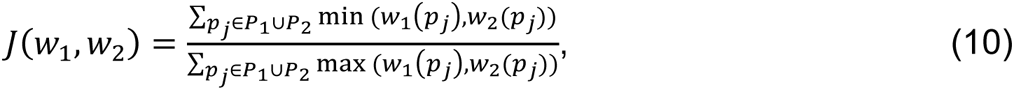

where *w*_1_ and *w*_2_ are the weighted spatial vectors of event 1 and event 2, and *P*_1_ ∪ *P*_2_ denotes the union set of pixel positions for these two events.

Once the crucial keys have been addressed, the clustering algorithm commences by treating each signal event as an individual group. It then iteratively clusters the most spatially consistent signal groups until the consistencies of all pairs are below the user-defined consistency. The consistency is set to 0.3 by default. In a subsequent step aimed at eliminating CFUs likely comprised of false positive events, an event number filter is applied, utilizing a default threshold of 3. For each remaining signal group, AQuA2 transforms the average spatial vector into a visual map to showcase the CFU’s position. Additionally, it captures the 50% rising times of the events within the group to construct the corresponding occurrence sequence for that CFU. These occurrence sequences play a pivotal role in quantifying the interactions between CFUs in subsequent analyses.

###### Settings for AQuA2 and its peer methods

In order to showcase the optimal performance of peer methods and ensure a fair comparison among them, we fine-tune their input parameters prior to application.

###### AQuA2

We apply the MATLAB version of AQuA2 to the synthetic data with the following parameters: We set the boundary removal “regMaskGap” to 0, minimum size “minSize” to 20, threshold “thrARScl” to 3, smoothing parameter “smoXY” factor to 1, baseline cut “cut” to 250, and minimum duration “minDur” to 2. All other parameters are set by default. In real-world applications, for optimal performance, fundamental parameters such as “minSize”, “minDur”, and “thrARScl” might be adjusted based on the specific characteristics of the data.

###### AQuA

We apply the MATLAB version of AQuA with the following parameters: We set “regMaskGap” to 0, “smoXY” to 1, “thrARScl” to 3, and “minSize” to 20. All other parameters are set by default. Notably, when comparing AQuA and AQuA2 in real applications, they shared the same basic parameter settings for a fair comparison.

###### Begonia

We apply the Begonia, a MATLAB package, to the synthetic data. The script code we used is the demo code for processing a single recording, with the following modifications: We let Begonia estimate the baseline using its default method rather than the 5th percentile of the first 100 frames. We set the min duration parameter “roa_min_duration” to 5, and the minimum size to be 20 to make it consistent with other event-based methods. The resultant activity regions (ROAs) are directly considered as events and compared with the ground truth.

###### CaImAn

We apply the newest Python version of CaImAn to the synthetic data. The script we use is modified based on “demos/notebooks/demo_pipeline.ipynb”. The changes are listed below: Since there is no registration issue, we bypass the motion correction part. With the known temporal template of the signals, we specified the decay speed “decay_time” to 0.3. According to the spatial templates in our ground truth, we set the expected half size of neurons “gSig” to 15×15 and the half size of the patches to 50. The number of components per patch “K” is set to 10, while merging threshold “merge_thr” is specified as 0.5. Additionally, the amount of overlap between patches “stride_cnmf” is set to 10. To compare with ground truths, we extract the ROIs “estimates.A”, their dFF “estimates.F_dff”, and their decomposed spikes “estimates.S” to form spatiotemporal events.

###### suite2p

We apply the newest Python version of suite2p to the synthetic data. Since there is no registration issue, we set the parameter “do_registration” to False. The decay speed “tau” is specified as 0.3. Moreover, considering the spatial template we use, we set “spatial_scale” to 1 for good performance. Three features, ROIs’ position “stat”, ROIs’ curves “F”, and decomposed spikes “spks” are used to form spatiotemporal events.

### QUANTIFICATION AND STATISTICAL ANALYSIS

#### Workstation configuration

All experiments were conducted on a workstation with Intel(R) Xeon(R) Gold 6140@2.30Hz processor, 128GB memory, running Windows 10 64-bit and using Microsoft VC++ compiler. No GPU was used. Two ROI-based methods were implemented in Python, while the other three event-based methods were implemented in MATLAB.

#### Event evaluation for ROI-based methods

AQuA2, AQuA, and Begonia are event-based methods that directly use the detected events as results, eliminating the need for further processing. In contrast, two ROI-based methods, CaImAn and suite2p, extract ROIs and their corresponding dF and decomposed spikes as results. Since ROIs are only spatial concepts, they cannot be used to compare with spatiotemporal ground truth events. Thus, we integrate spikes, dF, and ROIs into events for comparison.

To form spatiotemporal events based on ROIs, we first estimate the standard deviation of noise, denoted as *σ*, in the average dF curve of the target ROI. Time windows larger than 3*σ* are considered as candidate time windows that may contain signals. Next, we utilize the information from decomposed spikes. We apply a filter to the spikes to remove false positives and assume that each remaining spike represents one detected signal event. We use these spikes as the center of the signal to segment the candidate time windows temporally. By combining the spatial ROI and segmented time windows, we can obtain signal events for performance evaluation.

#### Metrics for performance evaluation

To assess the accuracy of different detection methods, we employ two metrics: the weighted Intersection over Union (wIoU) and the F1 score. The wIoU evaluates the quality of detection at the pixel level while the F1 score measures the match relationship between detections and ground truths at the signal level.

#### wIoU

Motivated by the intuition that the brighter pixels should be assigned with larger weight, we use weighted IoU rather than the standard IoU to evaluate the quality of detection. This approach also helps to eliminate the impact of ground truth selection. The wIoU for a pair of detection *D*_*i*_ and ground truth *G*_*j*_, (*D*_*i*_, *G*_*j*_), can be expressed as,

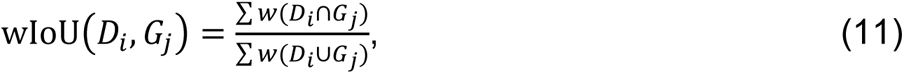

where the weight *w* of each pixel is the assigned weight according to the intensity.

The wIoU score ranges from 0 to 1, with a high score indicating a good match. However, pixels outside the ground truth are often of low intensity, which may result in an overestimation of the score if the detection includes many false positive pixels. For example, consider an extreme case where a detection *D*_*i*_ completely encompasses the corresponding ground truth *G*_*j*_, but also includes many pixels with 0 intensity such that its size is 10 times larger than that of *G*_*j*_. In this case, the estimated wIoU(*D*_*i*_, *G*_*j*_) will be 1, which is not reasonable. To avoid this issue, we set a lower bound of weight c = 0.1 that *w*(*pixel*_*k*_) = max (*c*, *I*(*pixel*_*k*_)) where *I*(*pixel*_*k*_ is the intensity of *pixel*_*k*_ in ground truth data, and *w*(*pixel*_*k*_) is the weight assigned to *pixel*_*k*_.

For each detected event *D*_*i*_, we define the highest score among all pairs between it and all ground-truth events as wIoU_*Di*_. Similarly, for each ground-truth event *G*_*j*_, the corresponding highest score is denoted as wIoU_*Gj*_. We can then calculate an overall score by averaging the score of both detections and ground truths using the following formula:

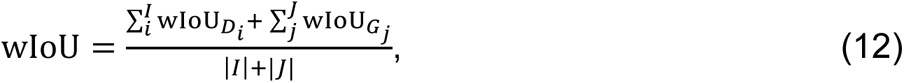

where *I* and *J* are the sets of indexes of detections and ground truths, respectively, and |*I*| and |*J*| are the number of indexes in *I* and *J*.

#### F1 score

A pair of detection *D*_*i*_ and ground truth *G*_*j*_ matches when the following condition holds:

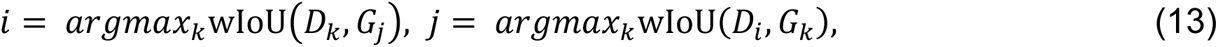

where D and G represent the detection set and ground truth set, respectively. That means that *D*_*i*_/*G*_*j*_ is the most closely matched detection/ground truth for *G*_*j*_/*D*_*i*_ among all other options at the pixel level. This definition helps to eliminate the impact of ground truth selection, as the match relationship remains unchanged regardless of whether pixels higher than 20% or 50% are selected as ground truth.

Let the number of matches be denoted as TP, the number of detections with no corresponding ground truth as FP, and the number of ground truths with no corresponding detection as FN. The F1 score can then be calculated as follows:

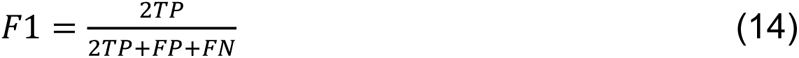

#### Interaction analysis of CFUs

Given that the curves of two correlated CFUs may have different waveforms, baseline trends, or contamination from signals not originating from the measured CFU due to potential overlap, directly calculating the correlation coefficient is not suitable. Instead, AQuA2 uses the occurrence sequences as features to analyze the interaction between two CFUs.

We propose a new statistical approach by examining the significance of co-occurrence between events in two sequences. Assuming the null hypothesis that each sequence follows a Poisson process with density *λ*, we can calculate the probability *p* that at least one event occurs within a time window of duration *L*.

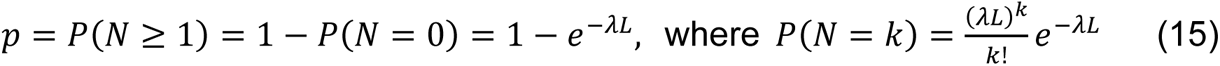

Taking one sequence, say sequence 2, with *n* events in it as a conditional sequence, we can place *n* time windows centered at the events in sequence 2 and count how many windows have at least one event from sequence 1. Note that the probability of a time window having at least one event is *p* under the null hypothesis. Assuming there are *m* windows that meet the criteria, meaning *m* co-occurring pairs of events in two sequences, the significance of dependency can be evaluated by,

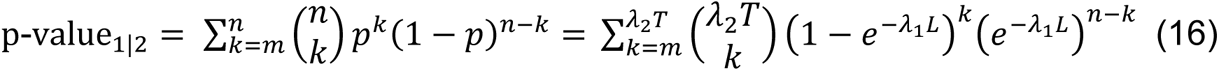

where *λ*_1_ and *λ*_2_ represent the densities of event sequences in CFU1 and CFU2, respectively. *L* is the window size that we may judge two events co-occur, and *T* denotes the sequence length. This is a test based on the Binomial distribution (n,m,p).

Notably, to avoid the situation where two events in one sequence refer to the same event in another sequence, our code does not allow time windows to overlap. This results in different lengths of time windows for different events. In this case, the saddle-point approximation method ^62^ is used to efficiently calculate the p-value.

During the analysis of the interaction between each pair of CFUs, AQuA2 will, by default, switch the conditional sequence and gradually adjust the length of the time window *L* until it reaches a user-defined value. The minimum p-value is then used as the final measure of significance to describe their dependency. It is important to highlight that the proposed methodology exhibits directionality, implying that we can take a specific sequence as a condition. For instance, the sequence of some CFU, or a stimulus sequence. Moreover, it doesn’t necessitate the time window to be centered around the event, permitting shifts in the windows.

#### The output features of AQuA2

The output features of AQuA2 can be divided into two categories: event-level features and CFU-level features. All the results can be obtained in the output “res.mat” file. Event-level features focus on the quantification of individual events, while CFU-level features describe the active functional units in the fluorescence imaging data.

Event-level features can be further divided into three classes: (1) Basic features of individual events, including voxel set, duration, area size, average curve of spatial footprint, average dF curve of spatial footprint, rising time, peak p-value, area under the curve (AUC), and others. These features serve as filters to select valuable signals and can also be utilized for statistical analysis, such as comparing changes in signal activity before and after the application of stimuli/drugs in control experiments. (2) Propagation-related features, including propagation speed, propagation map, and propagation trend in various directions. These describe the propagation information of events and can be used to uncover potential signal pathways. (3) Network features, which involve the distances between events and user-defined regions (e.g., cell regions or landmarks), as well as assessing the co-occurrence of events in spatial or temporal dimensions. The former reflects the association between specific regions and events, while the latter can be used to analyze the collective characteristics of events, such as signal bursts.

CFU-level features can also be categorized into three types: (1) Individual CFU information, including the spatial map, event sequence, average curve, and average dF curve. This is the result of CFU identification and can be utilized to visualize the active functional units in the data. (2) The dependency between every pair of CFUs, as well as the relative delay between two CFUs. This is the result of CFU interaction analysis. Users could utilize this information to understand the interaction between CFUs. (3) The information of CFU groups, including CFU indexes and the relative delay of each CFU. This is the result of further postprocessing, which employs hierarchical clustering based on the dependency between CFU pairs. It can cluster CFUs into groups, show their relative order, and may unveil circuit pathways.

## ACKNOWLEDGEMENTS

This work was primarily supported by the National Institutes of Health (NIH) grant U19NS123719 (G.Y., A.N.). It was partially supported by the NIH grants R01MH110504 (G.Y.), R01NS099254 (K.P.), R01MH121446 (K.P.), U19NS112959 (A.N.), NSF CAREER 1942360 (K.P.), the Howard Hughes Medical Institute (M.A.), the Sol Goldman Charitable Trust (A.N.), equipment funds from C. and L. Greenfield (A.N.), Wellcome Trust Senior Research Fellowship (214244/Z/18/Z, D.A.L.), Marie Skłodowska-Curie action “ZENITH” (H2020-MSCA-ITN-2018 813457, D.A.L.) and Elite network of Bavaria/ ENB Biological physics program award (P.N.B.). The content is solely the authors’ responsibility and does not necessarily represent the official views of the NIH.

## AUTHOR CONTRIBUTIONS

Conceptualization, X.M. and G.Y.; Methodology, X.M. and G.Y.; Software. X.M. and M.B.; Investigation, J.L., A.C., V.R., M.R., C.T. D.D., E.C., P.B., R.A., D.L., K.P., M.A., and A.N.; Writing – Original Draft, X.M.; Writing – Review & Editing, X.M., W.Z., Y.Z., C.T., A.N., M.A., K.P., and G.Y.; Supervision, G.Y.

## INCLUSION AND DIVERSITY

We support inclusive, diverse, and equitable conduct of research.

---

**Figure S1, related to Figure 1.**
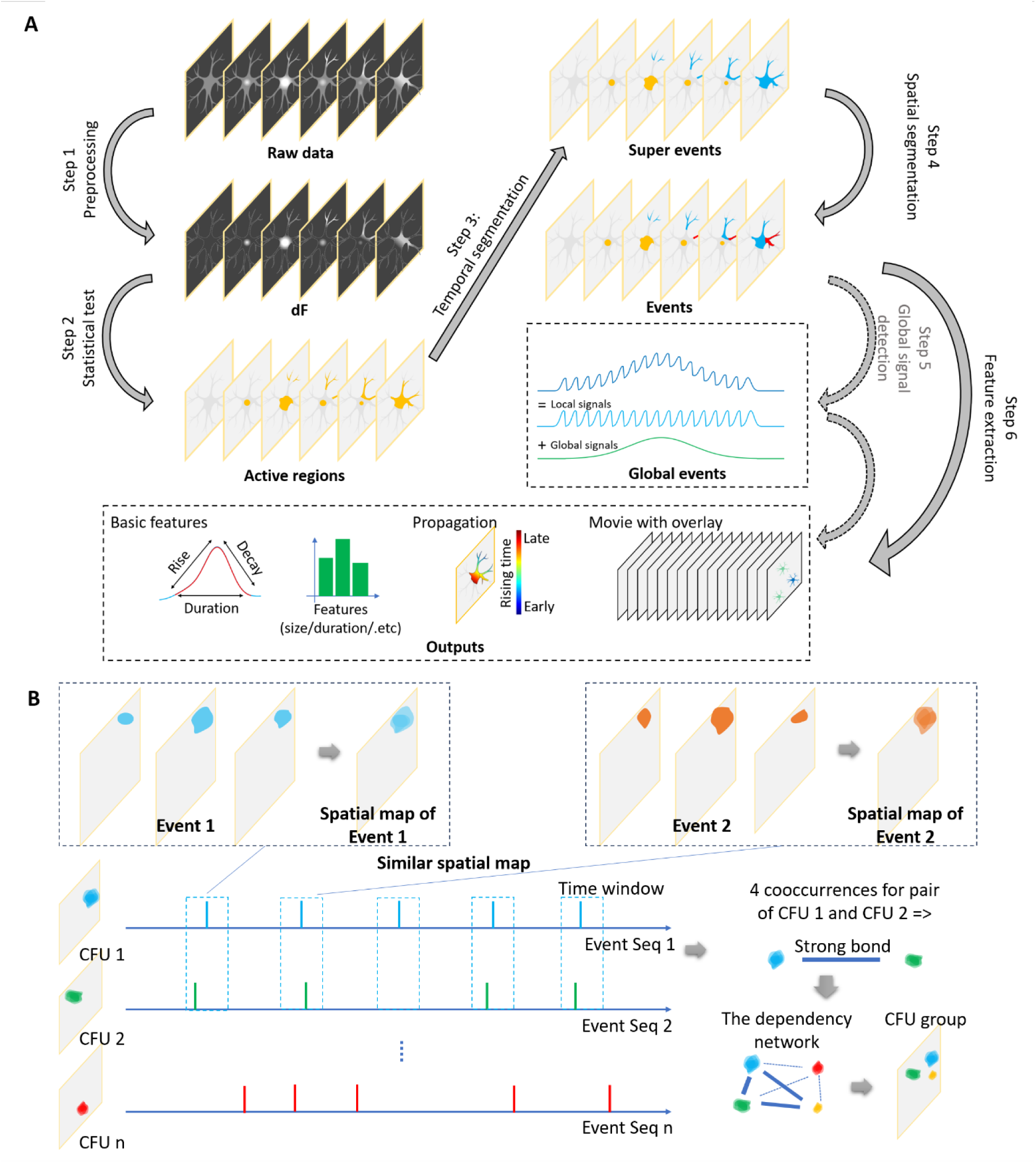
Principles of AQuA2. (A) In-depth pipeline providing a comprehensive overview of intermediate results, detailing every step of the event detection procedure in AQuA2: preprocessing, statistical testing, temporal segmentation, spatial segmentation, optional global signal detection, and final feature extraction. Each colored region corresponds to a distinct result in each step. (B) Details on the CFU module, beginning with the conversion from spatiotemporal events into spatial maps, followed by CFU interaction analysis and CFU grouping.

**Figure S2.**
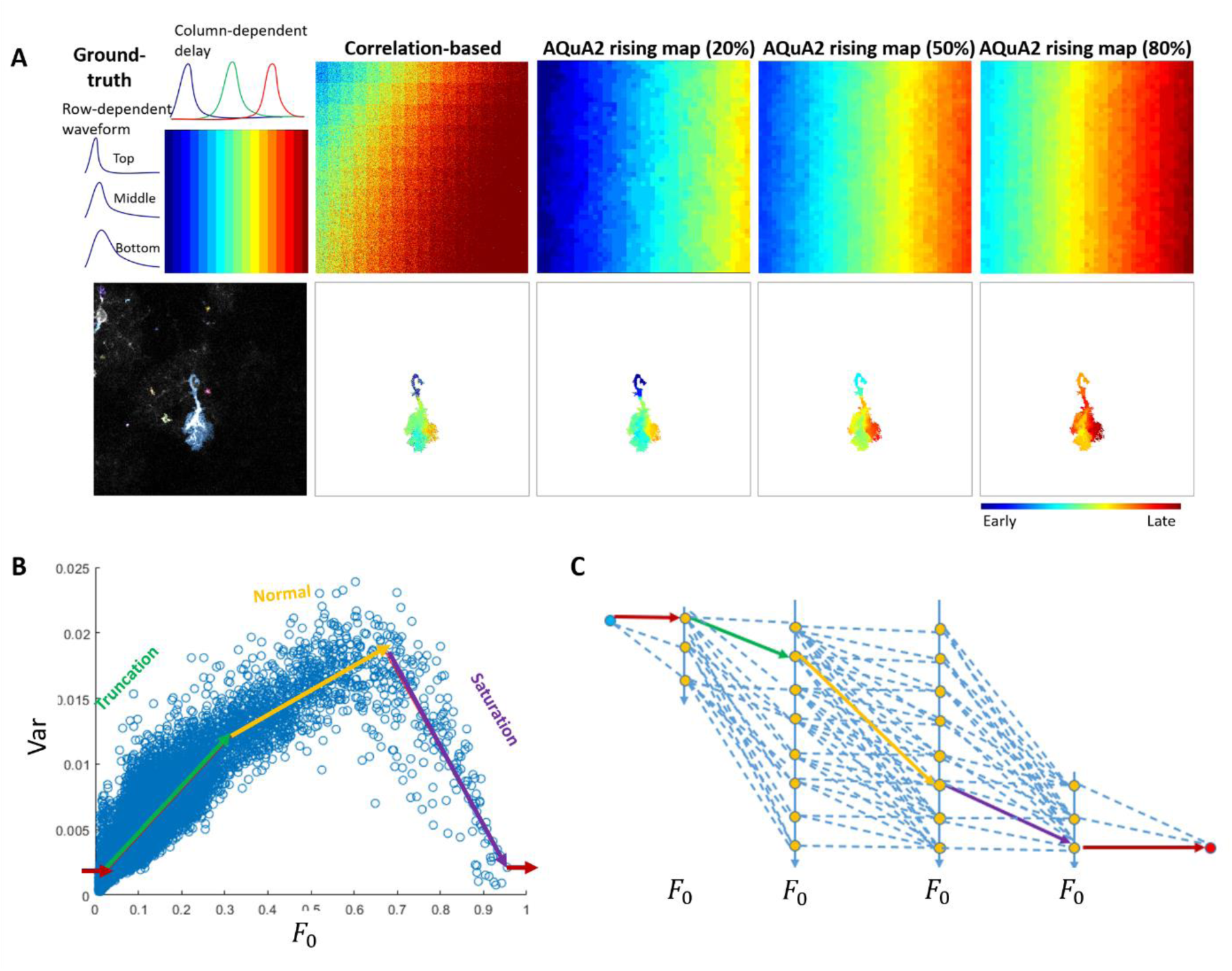
Supplementary diagrams for method parts. (A) A comparison between the correlation-based method and AQuA2 for propagation analysis. The top row displays simulation data featuring column-dependent delay and row-dependent waveform. The bottom row exhibits ex vivo astrocytic calcium recordings from mice. As demonstrated, the rising map generated by the correlation-based method tends to be noisy and prone to the influence of waveform, often leading to suboptimal outcomes. In contrast, AQuA2 employs joint alignment techniques, enabling it to deliver smooth, comprehensive, and precise information. (B) The noise variance versus baseline value of all the pixels. Each blue circle represents one pixel/sample. The green, yellow, and purple line segments of the piecewise linear function represent the relationships under truncation, normal, and saturation cases, respectively. The horizontal red line segments represent the noise variance for outliers. (C) The graph model for solving the piecewise linear function. Using the shortest path algorithm, the model could be solved efficiently. Each node on the vertical line represents the candidate ends of line segments, each edge represents the summation of absolute differences between pixels’ noise variance and the line segment linking two candidate ends. Thus, the cost of the shortest path is equal to the piecewise linear function with the minimum absolute error.

**Figure S3.**
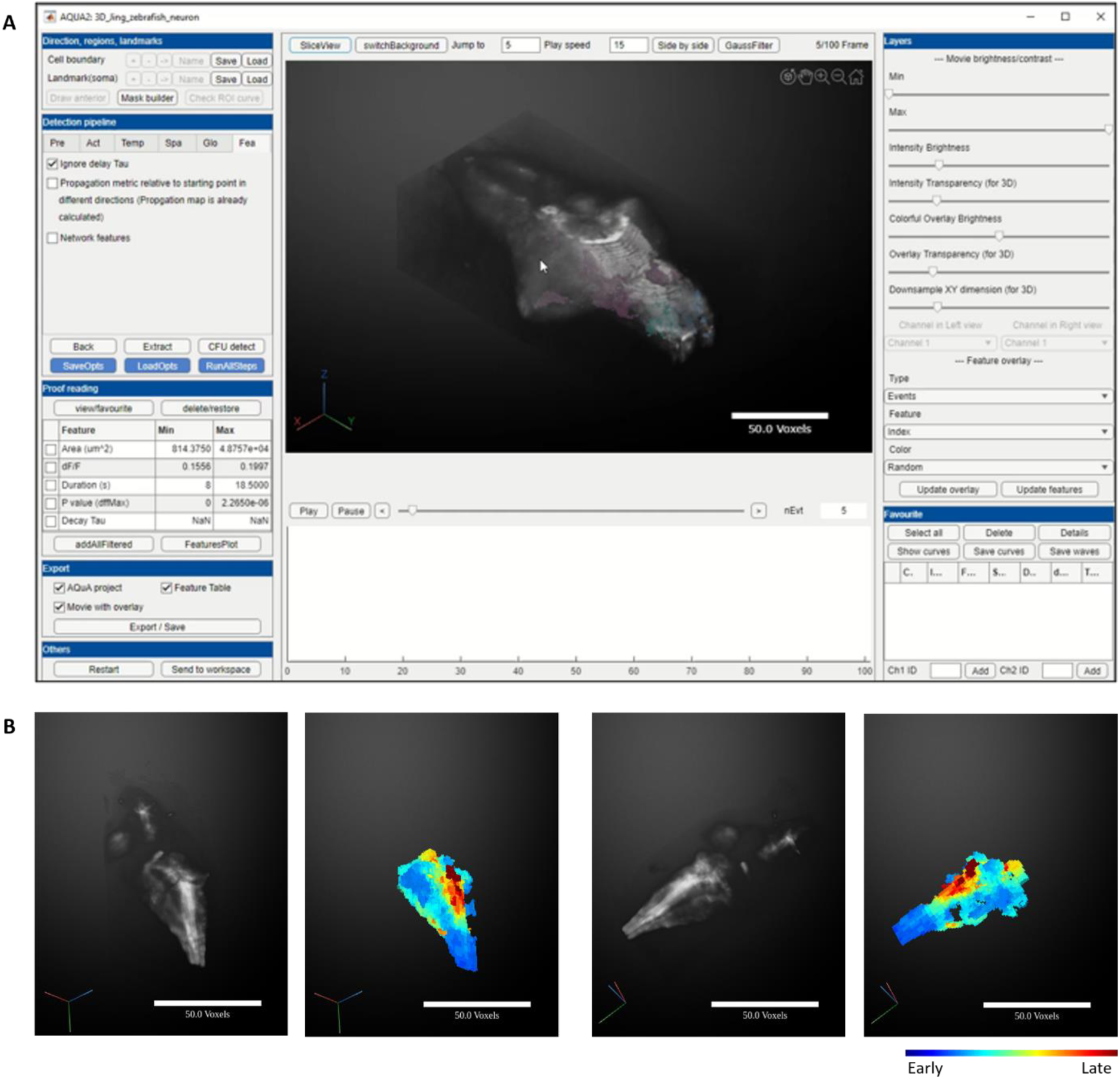
AQuA2 can be applied to 3D time-lapse imaging data. (A) AQuA2 GUI schematic diagram for 3D data. Each colored region represents one detected event. (B) The signal propagation pattern was detected by applying AQuA2 on 3D astrocytic calcium recording in zebrafish expressing *Tg(GFAP: jRGECO1B)* under caffeine influence, with blue indicating early rising time and red indicating late rising time.

**Figure S4.**
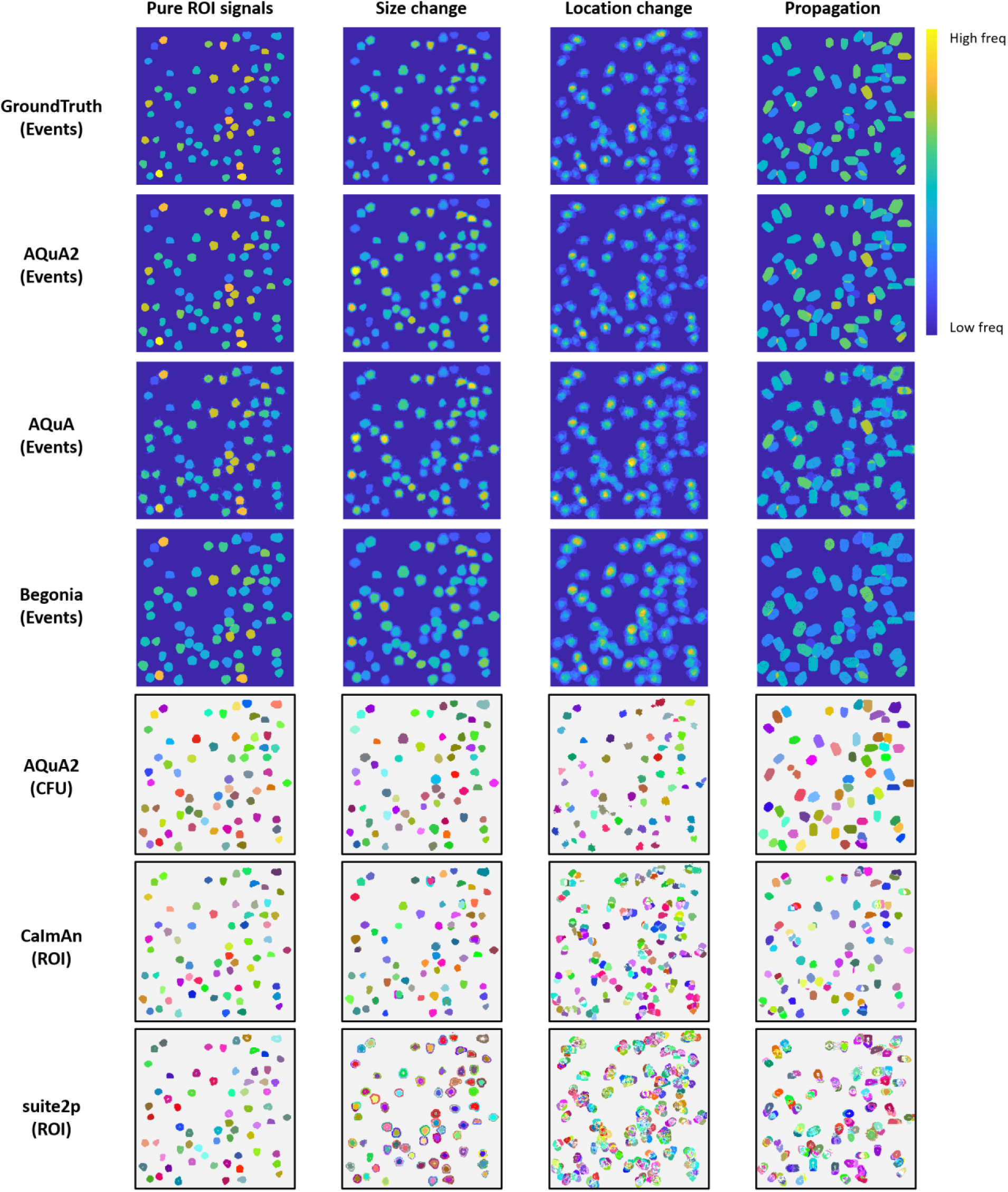
Detection results of peer methods on different experimental settings under 10dB SNR. From the first column to the fourth column, the experimental settings are pure ROI signals, signals with a size-change odd value of 2, signals with a location change value of 2, and signals with a propagation frame value of 6, respectively. For the ground truth and event-based methods, we use the color to represent the frequency of events at each pixel (with dark blue and bright yellow representing the lowest and highest frequencies, respectively). For the ROI-based methods, we visualized each detected ROI by a colored region and compared the results of CaImAn and suite2P with the CFU results of AQuA2.

**Figure S5.**
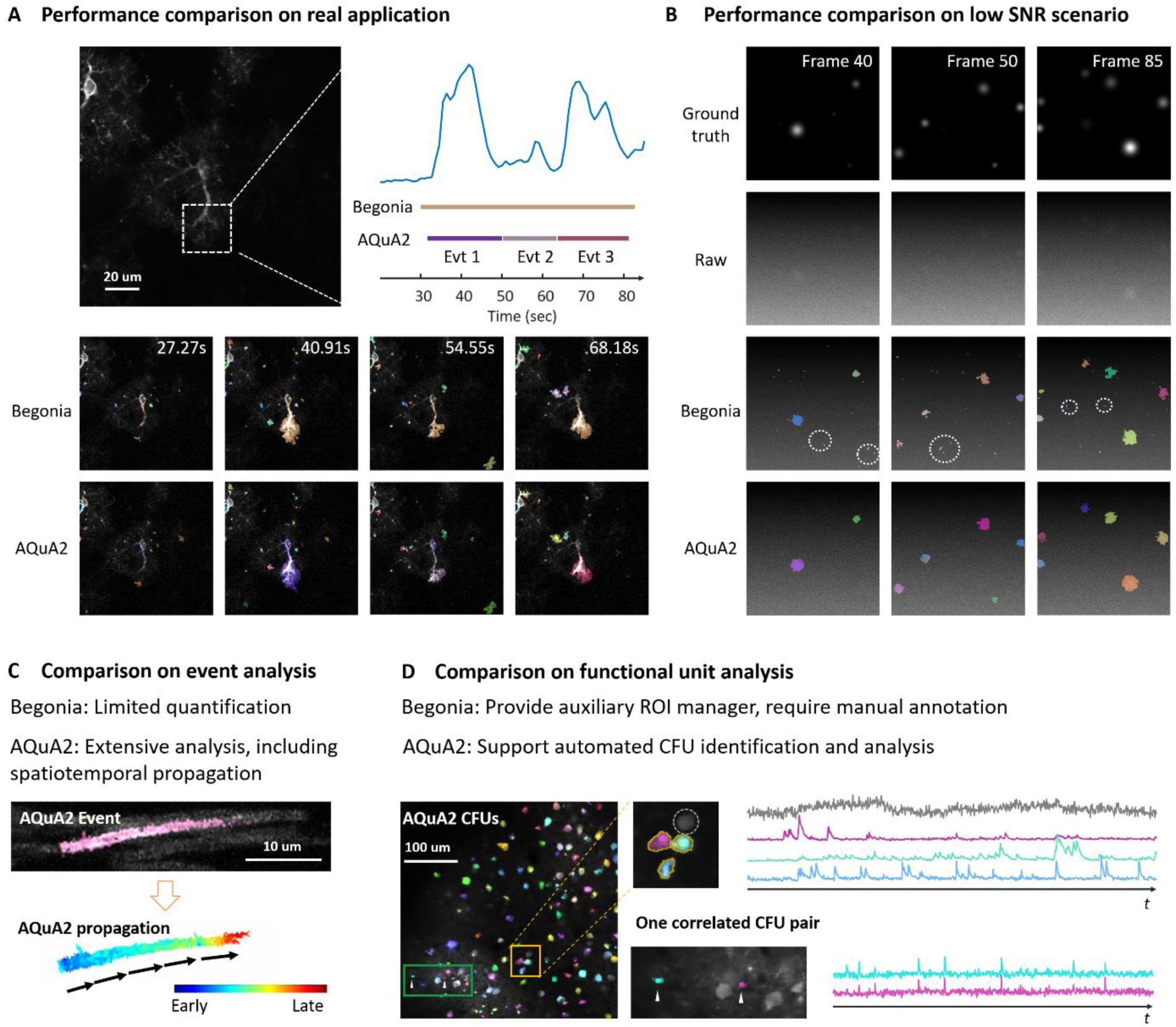
Comparison of Begonia and AQuA2. (A) Performance comparison on experimental data (*ex vivo* astrocytic calcium imaging in a mouse brain slice ^14^). Each colored region indicates one detected event. A white dashed box highlights the major signals of interest, and its average curve is shown on the right. The time intervals of detected signals by the two methods are marked with line segments in corresponding colors. (B) Performance comparison on synthetic data under low SNR. From top to bottom, the rows display ground truth, raw data, Begonia detection results, and AQuA2 detection results. Each colored region indicates one detected event. White dashed circles mark example false positives. (C) Comparison of event analysis. Begonia allows limited quantification, while AQuA2 supports extensive analyses, including propagation quantification. An example of propagation analysis (oligodendrocyte myelin sheath calcium imaging data from *Tg(mbp:memGCaMP7s)* zebrafish) is shown, with blue indicating early rising times and red indicating late rising times. (D) Comparison of functional unit analysis. Begonia only provides an auxiliary ROI manager and requires manual annotation by the user, while AQuA2 supports automated CFU identification and analysis. An example of AQuA2’s CFU module (neuronal calcium imaging data from the mouse visual cortex) is shown. Each colored region indicates one detected CFU. Although some neurons are visible in the background, they are silent in the experiment and thus not detected.

**Figure S6.**
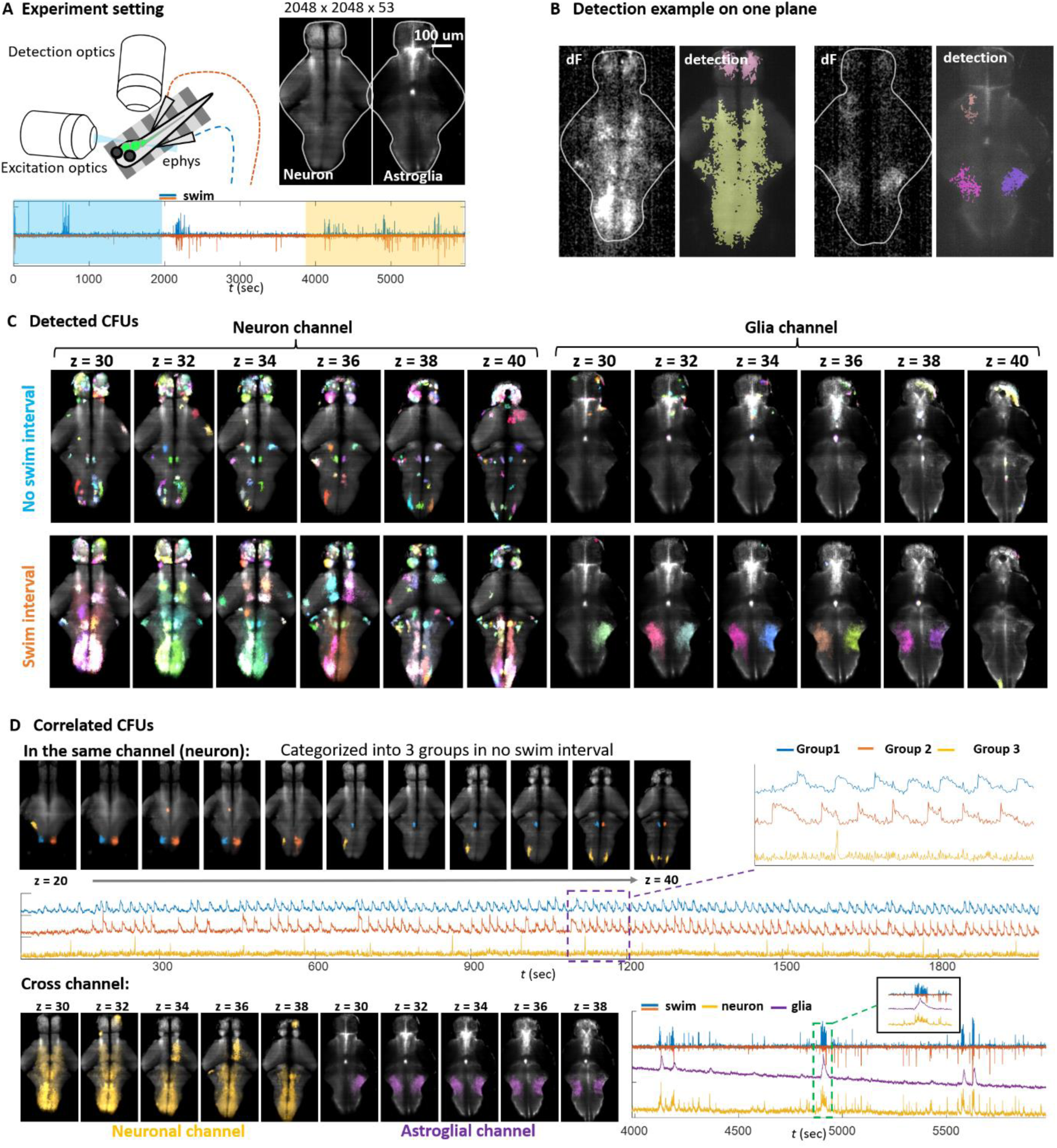
Application of AQuA2 on dual-color zebrafish recording. (A) Virtual environment for fictively swimming paralyzed fish. A custom light sheet microscope was used to record neuronal calcium (*Tg(ELAVL3: GCaMP8f)*) and astroglial calcium (*Tg(GFAP: jRGECO)*). The frame rate is 3.01 Hz. Swim commands were detected by two probes. During the whole duration, two time windows were selected, labeled by blue and yellow masks, to observe the zebrafish signals in the non-swim interval or swim interval. (B) Detection example on one plane. Each colored region represents one detected signal event. (C) Detected CFUs of two intervals on different planes. Each colored region is one CFU. (D) Obtained correlated CFUs. Top: three different CFU groups on different planes in non-swim intervals, with their average curves shown below. Bottom: Correlated CFUs found in the cross channel of the swim interval with their average curves shown on the right.

**Figure S7.**
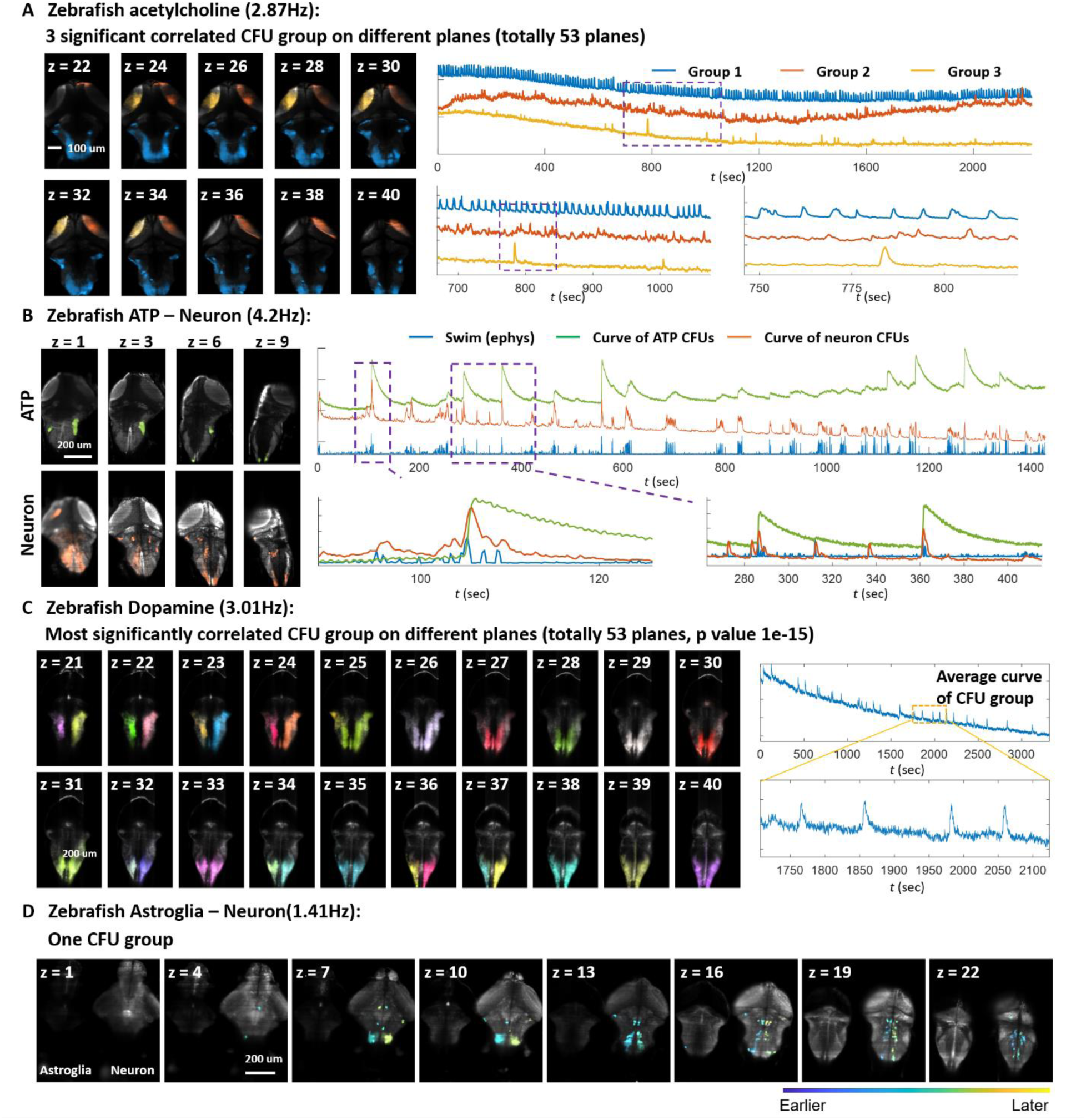
Supplementary applications of AQuA2. (A) Application of AQuA2 to identifying CFU groups on the zebrafish acetylcholine imaging. During the experiment, paralyzed zebrafish performed fictive swimming in a virtual reality environment. Acetylcholine signals were expressed using *Tg(ELAVL3: AChSnFR)* and captured using a custom light sheet microscope with a z-resolution of 5um/px and x and y resolutions of 0.40625um/px. The frame rate was set at 2.87 Hz. Each CFU group is labeled with a distinct color, with their average curve shown on the right. Each group shows a distinct signal pattern. (B) Utilizing AQuA2 to identify correlated CFUs between the ATP channel and Neuron channel in zebrafish, despite waveform differences. ATP and neuronal signals were expressed using *Tg(ELAVL3:ATP)* and *Tg(ELAVL3:jRGECO)*, respectively, recorded by a custom light sheet microscope with a z-resolution of 10um/px and x-y resolutions of 0.40625um/px. The frame rate was set at 4.2 Hz. The average curves of CFUs for both channels (green and orange) are displayed on the right, alongside swim strength (blue). Zoom-ins of two time windows are provided. The obtained CFUs demonstrate a strong correlation with swimming behavior. (C) Application of AQuA2 to identifying the most significant CFU group on different planes of zebrafish dopamine imaging. Each colored region represents one detected CFU, with the average curve of all these CFUs illustrated on the right. The dopamine was expressed using *Tg(GFAP: GRAB-DA)* and recorded by a custom light sheet microscope with a z-resolution of 4.9um/px and x and y resolutions of 0.40625um/px. The frame rate was set at 3.01 Hz. This application demonstrates AQuA2’s capability to quantify dopamine activity. (D) Application of AQuA2 to the CFU group identification and analysis in astroglia-neuron imaging in zebrafish. Astroglial calcium and neuronal calcium were expressed through *Tg(ELAVL3: GCaMP7f; GFAP: jRGECO1B)*. Each connected colored region is one CFU, and the color shows relative delay in this group. Using AQuA2, potential brain circuits can be found through CFU grouping.

